# PtdIns(3,4)P_2,_ Lamellipodin, and VASP coordinate cytoskeletal remodeling during phagocytic cup formation in macrophages

**DOI:** 10.1101/2022.03.08.483476

**Authors:** Fernando Montaño-Rendón, Glenn F. W. Walpole, Matthias Krause, Gerald R.V. Hammond, Sergio Grinstein, Gregory D. Fairn

## Abstract

Phosphoinositides are pivotal regulators of vesicular traffic and signaling during phagocytosis. Phagosome formation, the initial step of the process, is characterized by local membrane remodelling and reorganization of the actin cytoskeleton that leads to formation of the pseudopods that drive particle engulfment. Using genetically-encoded fluorescent probes we found that upon particle engagement a localized pool of PtdIns(3,4)P_2_ is generated by the sequential activities of class I phosphoinositide 3-kinases and phosphoinositide 5-phosphatases. Depletion of the enzymes responsible for this locally generated pool of PtdIns(3,4)P_2_ blocks pseudopod progression and ultimately phagocytosis. We show that the PtdIns(3,4)P_2_ effector Lamellipodin (Lpd) is recruited to nascent phagosomes by PtdIns(3,4)P_2_. Furthermore, we show that silencing of Lpd inhibits phagocytosis and produces aberrant pseudopodia with disorganized actin filaments. Lastly, vasodilator-stimulated phosphoprotein (VASP) was identified as a key actin-regulatory protein mediating phagosome formation downstream of Lpd. Mechanistically, our findings imply that a pathway involving PtdIns(3,4)P_2_, Lpd and VASP mediates phagocytosis at the stage of particle engulfment.

## Introduction

Phagocytosis, the process whereby cells engulf and dispose of effete cells, microorganisms, and foreign particles, is pivotal for immunity and tissue homeostasis^1, 2^. Phagosome formation, the initial step during phagocytosis, entails marked reorganization of the actin cytoskeleton and membrane remodelling events that lead to pseudopod extension and particle engulfment^3, 4^. Earlier studies have suggested that phosphoinositides are pivotal molecules in the regulation of phagocytosis. Phosphoinositides, which play crucial roles in signaling and membrane traffic^5^, reside primarily in the cytosolic leaflet of organelles including the plasma membrane and phagosomes. Phosphatidylinositol 4,5-*bis*phosphate (PtdIns(4,5)P_2_) is enriched in the plasma membrane where, amongst its many roles, it acts as a positive regulator of actin polymerization^6^. The conversion of PtdIns(4,5)P_2_ to phosphatidylinositol 3,4,5-*tris*phosphate (PtdIns(3,4,5)P_3_) is a potent stimulus resulting in alterations in actin dynamics through the activation/inactivation of Rho-GTPases, as well as promoting the hydrolysis of PtdIns(4,5)P_2_ by activating phospholipase Cγ. Indeed, it has been appreciated for nearly two decades that IgG-opsonized particle engagement by the phagocytic Fcγ receptor leads to the activation of the phosphoinositide-3-kinase (PI3K)^7^ that is required for the internalization of large phagocytic prey^8, 9^. Remarkably, the dynamic changes in phosphoinositides and the actin cytoskeleton during phagocytosis are restricted to the site of particle engagement and do not propagate to the rest of the plasma membrane. Accordingly, phagosome formation is characterized by the focal generation of PtdIns(3,4,5)P_3_^10^ at the site of contact and extending pseudopods, while a concomitant disappearance of PtdIns(4,5)P_2_ is observed at the base of the phagocytic cup prior to sealing of the nascent phagosome^11^.

The downstream effects of PI3K activation during phagocytosis have been mostly attributed to the generation of PtdIns(3,4,5)P_3_. However, some of these findings should be interpreted with caution, as most were obtained using probes with dual specificity for PtdIns(3,4,5)P_3_ and PtdIns(3,4)P_2_, such as the pleckstrin-homology (PH) domain of Akt^12, 13^. Recent advances in lipid bio-sensor design have enabled the distinction of the differential roles of PtdIns(3,4,5)P3 and the related PtdIns(3,4)P2^14, 15^. PtdIns(3,4)P2 is increasingly appreciated as involved in cellular processes such as endocytosis, macropinocytosis, cell migration, and neurite initiation^16,17,18,19,20^. To date, however, the role of PtdIns(3,4)P_2_ and its effectors during phagocytosis remains largely unknown. Using recently developed specific probes we demonstrate here that PtdIns(3,4)P_2_ accumulates greatly at sites of phagocytosis, where it persists long after PtdIns(3,4,5)P_3_ disappears. More importantly, we show that selective depletion of PtdIns(3,4)P_2_ attenuates particle internalization, highlighting the importance of this unique phosphoinositide.

## Results

### Detection of PtdIns(3,4)P_2_ in the plasma membrane of resting macrophages

The distribution of PtdIns(3,4)P_2_ in unstimulated RAW 264.7 macrophages was investigated first. To this end we employed a recently described genetically-encoded biosensor based on tandem carboxy-terminal PH (cPH) domains of TAPP1^21, 22^ . Using these biosensors, NES-mCh- cPHx3 and NES-EGFP-cPHx2 (cPHx3 and cPHx2 respectively), we detected a discrete plasmalemmal pool of PtdIns(3,4)P_2_ in resting cells (Figure 1A, left panel). To confirm the specificity and responsiveness of the probes we co-expressed inositol polyphosphate 4- phosphatase type II (INPP4B), which selectively hydrolyzes the D-4 position of PtdIns(3,4)P_2_ ^23, 24^ To ensure plasmalemmal targeting of INPP4B, the phosphatase was attached to a carboxy-terminal CAAX motif (INPP4B-CAAX) that is prenylated and polycationic. As shown in Figure 1A (middle and right panels) the biosensors detached from the membrane upon co-expression of INPP4B-CAAX, but not when the catalytically inactive INPP4B(C842A)-CAAX mutant was co-expressed. These observations validate the selectivity of the probes and confirm that modest yet detectable amounts of PtdIns(3,4)P_2_ are indeed present in the membrane of resting macrophages. Furthermore, we documented that production of PtdIns(3,4)P_2_ is PI3K- dependent, since treatment with nanomolar concentrations of the PI3K inhibitor wortmannin completely released the cPHx3 probe from the plasma membrane (Figure 1B,C).

**Figure 1.**
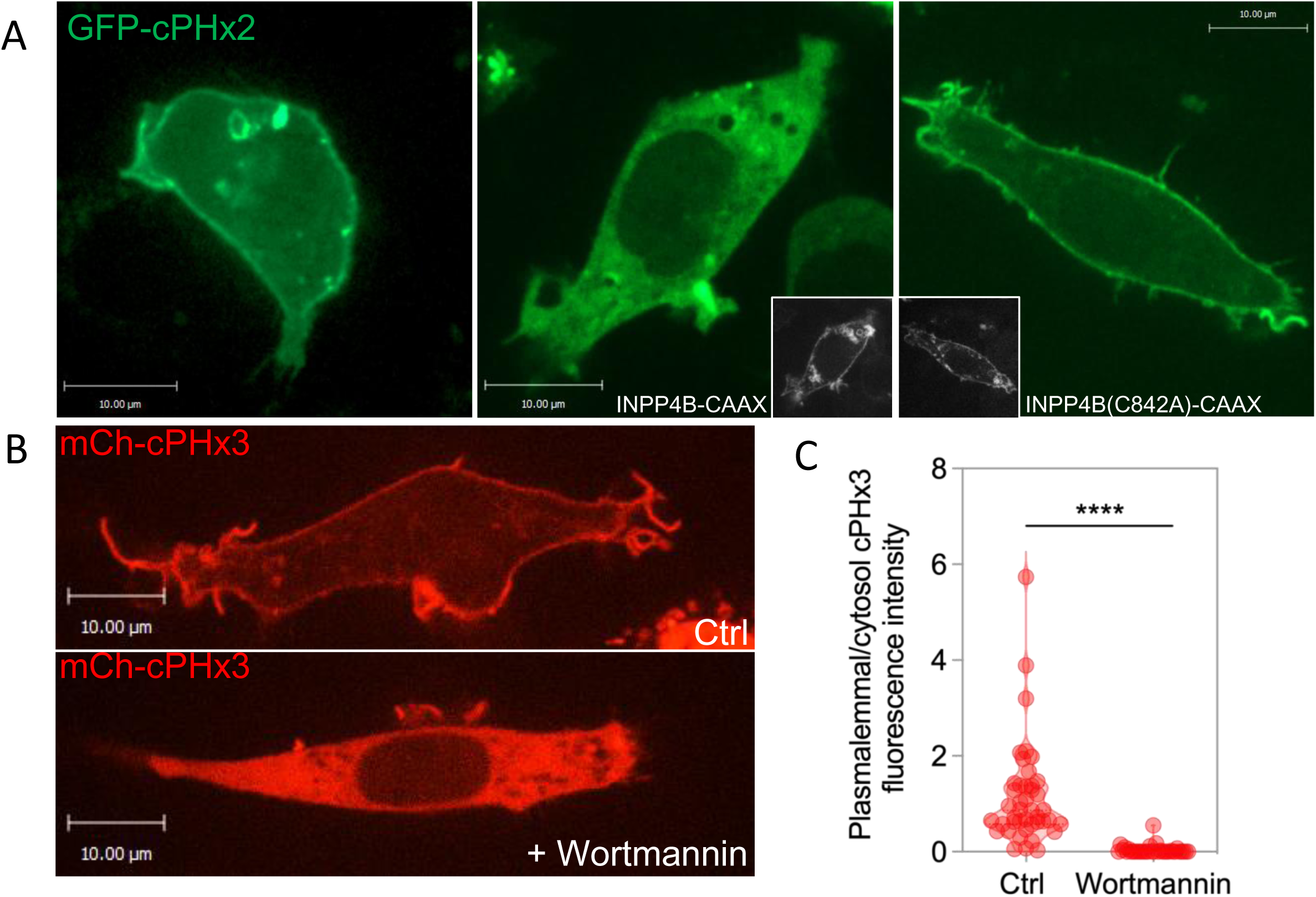
PtdIns(3,4)P_2_ at the plasma membrane of RAW 264.7 macrophages. (A) Confocal section of RAW 264.7 macrophages expressing GFP-cPHx2 alone (left) or in combination with either BFP-INPP4B-CAAX (middle) or BFP-INPP4B(C842A)-CAAX (right); insets represent the BFP. (B) Representative confocal micrographs of RAW 264.7 cells expressing mCherry-cPHx3 in either control (top) or 100 nM wortmannin-treated (bottom) conditions. (C) Normalized plasmalemmal mCherry-cPHx3 fluorescence intensity in control and wortmannin-treated cells. For quantitation, plasmalemmal fluorescence intensity was normalized to cytosolic fluorescence intensity. Unpaired *t*-test of individual values across n=3 independent experiments.

### PtdIns(3,4)P_2_ accumulates at the site of particle engagement during phagocytosis

Next, we examined the distribution of PtdIns(3,4)P_2_ during phagocytosis. Upon exposure to IgG-opsonized sheep erythrocytes (SRBCs), RAW 264.7 cells exhibited a marked accumulation of the PtdIns(3,4)P_2_ probe at the site of phagocytosis to levels greatly surpassing those of the neighbouring unengaged plasma membrane, an observation consistent with stimulated local production of the lipid (Figure 2A, left panel). PtdIns(3,4)P_2_ accumulated in the phagosomal cup and remained in the membrane even after scission of the nascent phagosome (Video S1.). Of note, there were marked differences in the distribution and dynamics of recruitment of cPHx2 and PH-BTKx2 (a sensor for PtdIns(3,4,5)P_3_) during phagocytosis. Specifically, PtdIns(3,4,5)P_3_ accumulated primarily – albeit transiently – at the base of the phagocytic cup, whereas PtdIns(3,4,)P_2_ was acquired slightly later, initially attaining higher levels at the tips of the extending pseudopods (Figure S1., Video S2.) and later persisting for a few minutes in the sealed phagosome. The accumulation of the PtdIns(3,4)P_2_ probe at the phagosomal cup was largely obliterated by co-expression of the INPP4B-CAAX construct, but no depletion was observed when the catalytically-inactive INPP4B(C842A)-CAAX construct was used as negative control (Figure 2A and B). These findings imply that PtdIns(3,4)P_2_ is produced locally at sites of phagocytosis.

**Figure 2.**
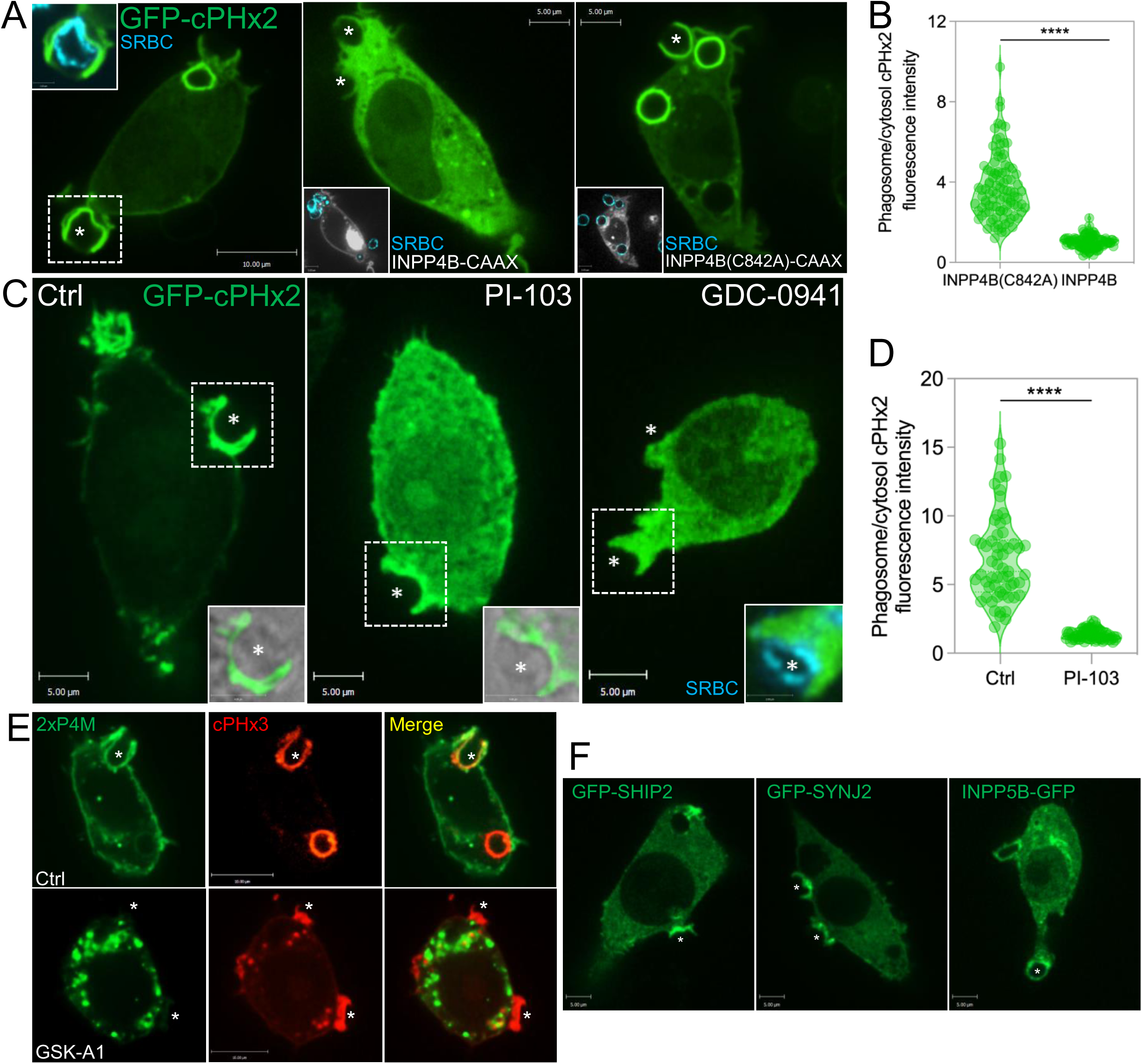
PtdIns(3,4)P_2_ accumulates at the site of particle engagement during phagocytosis, is susceptible to the degradation by INPP4B phosphatases, and its production is downstream of class I PI3Ks and 5’ phosphatases. (A) Confocal sections of RAW 264.7 macrophages expressing GFP-cPHx2 alone (left) or in combination with either BFP-INPP4B-CAAX (grey inset) (middle) or BFP-INPP4B(C842A)-CAAX (grey inset) (right) during phagocytosis. Asterisks represent sites of phagocytic cup formation. SRBCs are shown in blue. (B) Normalized phagosomal-cPHx2 fluorescence intensity in INPP4B(C842A)-CAAX and INPP4B-CAAX. For quantitation, here and elsewhere, phagosomal fluorescence intensity was normalized to plasmalemmal fluorescence intensity. Unpaired *t*-test of individual values across n=3 independent experiments, data are mean ± SD. (C) Representative confocal micrographs of RAW 264.7 macrophages expressing GFP-cPHx2 and pretreated with either vehicle control (left), 500 nM PI-103 (middle), or 500 nM GDC-0941 (right). Asterisks denote particles. Insets represent the area denoted by the dotted box showing SRBCs. (D) Normalized phagosomal GFP-cPHx2 fluorescence intensity in control and PI-103- treated cells. Unpaired *t*-test of individual values across n=3 independent experiments, data are mean ± SD. (E) Representative confocal micrographs of RAW 264.7 macrophages co-expressing GFP-2xP4M and mCherry-cPHx3 during phagocytosis in either control (top) or GSK-A1-treated (bottom) conditions. Asterisks indicate sites of particle binding. (F) Representative confocal sections of RAW 264.7 macrophages expressing SHIP2 (left), SYNJ2 (middle), and INPP5B (right) phosphatases during phagocytosis. Asterisks indicate sites of particle binding.

### Phagosomal PtdIns(3,4)P2 is produced downstream of Class I PI3K and 5-phosphatases

We next sought to elucidate the pathway(s) responsible for the biosynthesis of PtdIns(3,4)P_2_ during phagocytosis. We considered two possibilities, which are not mutually exclusive: that the PtdIns(3,4)P_2_ was being produced by dephosphorylation of PtdIns(3,4,5)P_3_ and/or that it was synthesized from PI(4)P by class II PI3Ks. Class I PI3K is known to be activated during phagocytosis and its inhibition arrests engulfment by preventing full extension of the pseudopods around the particle^8, 9^. To assess its involvement, we treated cells with nanomolar concentrations of the pan-PI3K inhibitor PI-103 or the class I PI3K-selective inhibitor GDC-0941 and monitored PtdIns(3,4)P_2_ formation during phagocytosis (Figure 2C and D). In both cases the recruitment of cPHx2 was virtually eliminated, consistent with a major role for class I PI3Ks in the generation of PtdIns(3,4)P_2_. Nevertheless, we examined the possible contribution of Class II PI3Ks. For this, we acutely depleted the plasmalemmal pool of PtdIns(4)P using GSK-A1, a potent and specific inhibitor of the type III phosphatidylinositol 4-kinase PI4KA (PI4KIIIα) that is largely responsible for the generation and maintenance of plasmalemmal PtdIns(4)P; it is noteworthy that selective inhibition of PI4KIIIα by GSK-A1 does not acutely alter the plasmalemmal levels PtdIns(4,5)P_2_^25^, the substrate of class I PI3Ks. Upon treatment with GSK- A1, the recruitment of mCherry-cPHx3 to the phagocytic cup remained largely unaffected despite a profound depletion of PtdIns(4)P monitored by the high-avidity biosensor 2xP4M (Figure 2E). These findings strongly suggest that dephosphorylation of PtdIns(3,4,5)P_3_ is the main source of PtdIns(3,4)P_2_ production at the phagocytic cup.

The involvement of PI3Ks that generate PtdIns(3,4,5)P_3_ suggests that phosphoinositide 5- phosphatases are also required for the synthesis of PtdIns(3,4)P_2_ during phagocytosis. Multiple 5-phosphatases could in theory play a role in the production of PtdIns(3,4)P_2_. For instance, Src homology 2 (SH2) domain-containing inositol-5-phosphatase 2 (SHIP2) is involved in the production of PtdIns(3,4)P_2_ at sites of endocytosis^26^ and at invadopodia^27^. Importantly, SHIP2 has also been reported to translocate to sites of phagocytosis^28^, making it an attractive candidate to mediate the production of PtdIns(3,4)P_2_. However, overexpression of the catalytically-inactive mutant of SHIP2 (D607A), which is expected to exert a dominant-negative effect, had no effect over phagosomal levels of PtdIns(3,4)P2 (Figure S2 A and B). Similar negative results were observed when the cells were treated with either a SHIP2-selective inhibitor (AS1949490) or with the pan-SHIP1/2 inhibitor (K118) (Figure S2 C). We therefore turned our attention to other 5-phosphatases: there is compelling evidence that SHIP1^29^, INPP5E^30^, and OCRL^31^ are all recruited to nascent phagosomes. In addition, we identified synaptojanin-2 (SYNJ2) and INPP5B as being present at the site of phagocytosis (Figure 2F). It is therefore conceivable that multiple 5-phosphatases collaborate to dephosphorylate PtdIns(3,4,5)P_3_ to PtdIns(3,4,)P ^32^. Because of the likelihood of functional redundance, the identity of the specific phosphatases involved was not pursued further.

### Depletion of PtdIns(3,4)P_2_ impairs phagosome formation

We next sought to determine whether PtdIns(3,4)P_2_ accumulation is necessary for efficient FcγR-mediated particle uptake. To this end, phagocytic efficiency was compared in RAW 264.7 macrophages over-expressing INPP4B-CAAX to deplete plasmalemmal PtdIns(3,4)P_2_, or the catalytically inactive version INPP4B(C842A)-CAAX, used as a negative control. We observed a stark decrease in the phagocytic efficiency of cells expressing the INPP4B-CAAX, compared to cells expressing the INPP4B(C842A)-CAAX mutant (Figure 3A,B). In addition to the decrease in efficiency of internalization we noted a modest reduction in the number of SRBCs that were contacted by the INPP4B-CAAX expressing cells (Figure S4; Figure 3A, right panel).

**Figure 3.**
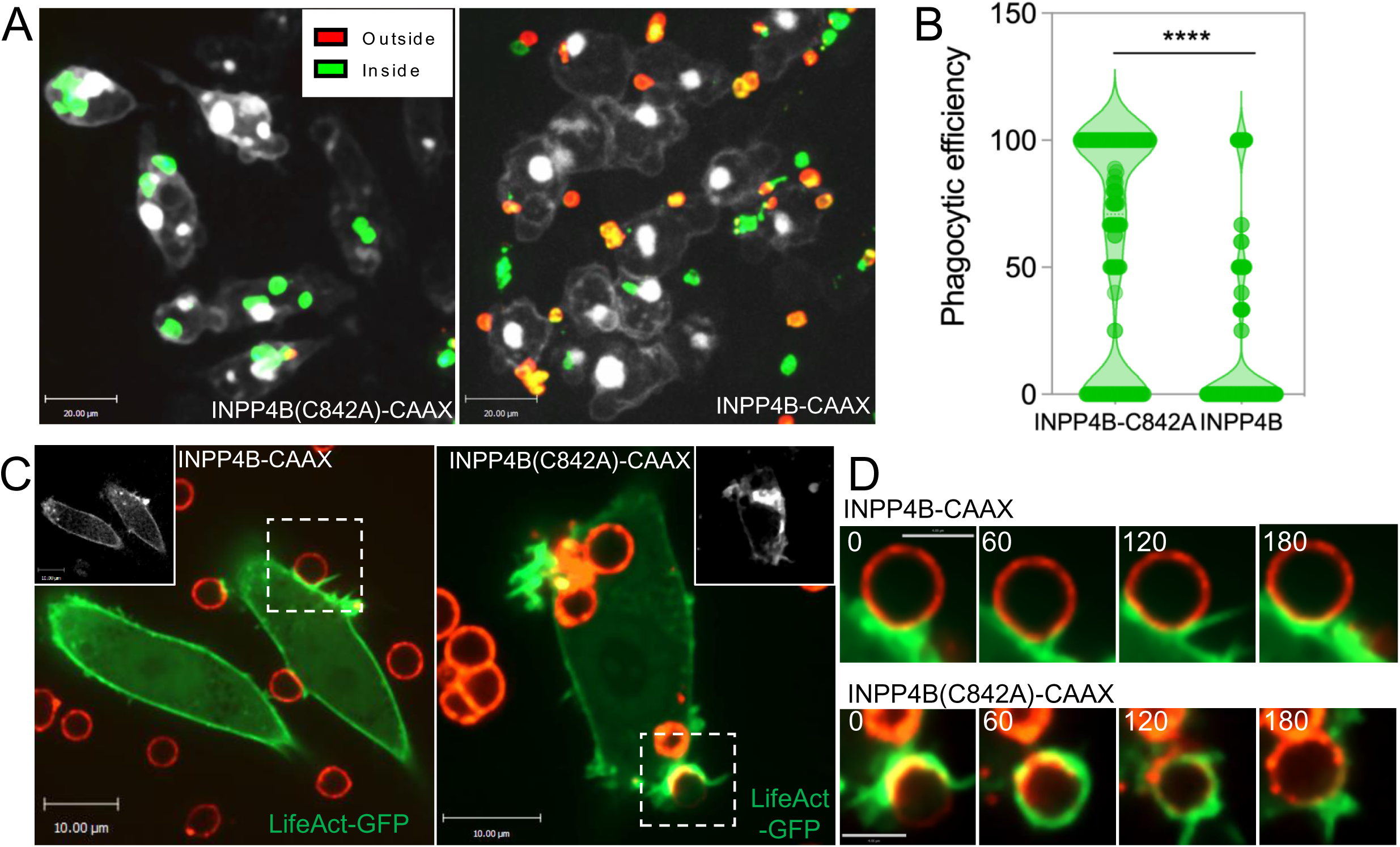
INPP4B-mediated depletion of PtdIns(3,4)P_2_ disrupts phagocytosis. (A) Representative confocal extended-focus projections of doxycycline-inducible RAW264.7 macrophages expressing BFP-INPP4B(C842A)-CAAX (Control) or BFP-INPP4B-CAAX (PtdIns(3,4)P_2_-depleted) following a 10 min incubation with IgG-opsonized SRBCs. Inside- outside staining was performed to differentiate between internalized/total (green) and non- internalized (red) SRBCs (see methods). (B) Quantification of the phagocytic efficiency in BFP- INPP4B(C842A)-CAAX and BFP-INPP4B-CAAX-expressing RAW 264.7 macrophages. Unpaired *t*- test of individual phagocytic efficiency values of n=228 and 267 cells, respectively across N=3 independent experiments; data are mean ± SEM. (C) Representative live confocal sections of RAW 264.7 macrophages expressing LifeAct-GFP and either BFP-INPP4B(C842A)-CAAX or BFP- INPP4B-CAAX during phagocytosis of IgG-opsonized SRBCs (red). Dotted boxes represent the phagocytic cups tracked in D. (D) Time-lapse microscopy of F-actin dynamics monitored by GFP- Lifeact during phagocytosis of IgG-opsonized SRBCs in RAW 264.7 macrophages expressing BFP- INPP4B-CAAX (top) or BFP-INPP4B(C842A)-CAAX (bottom) (0, 60, 120, and 180 sec timepoints). Scale bar = 3 μm. Similar results were observed in 10 independent experimental repeats.

Thus, we hypothesised that a defect in pseudopod extension or sealing must be responsible for the observed decrease in phagocytic efficiency.

Actin remodelling, which drives pseudopod extension, is a hallmark of the early stages of the phagocytic process. To assess the role of PtdIns(3,4)P_2_ in actin polymerization we performed live-cell imaging of cells co-expressing LifeAct-GFP and either INPP4B-CAAX or INPP4B(C842A)- CAAX. In cells expressing the inactive phosphatase, focal actin polymerization was followed by pseudopod formation, and progressive particle engulfment in a zipper-like fashion (Figure 3C, right; Fig. 3D, bottom row, and Video S2.). In contrast, cells where PtdIns(3,4)P_2_ was depleted by expression of INPP4B-CAAX, formed pseudopods that failed to progress and wrap around the phagocytic target, despite exhibiting an apparently normal initial actin polymerization (Figure 3C left; Fig. 3D, top row, and Video S3.). These findings indicate that PtdIns(3,4)P2 is necessary for phagocytosis and that it plays a direct role in regulating actin organization and pseudopod progression during the early stages of particle engulfment.

### Lamellipodin, a PtdIns(3,4)P_2_ effector, accumulates at the phagocytic cup

To date, only a handful of proteins have been identified as specific effectors of PtdIns(3,4)P_2_^18^. These include the modular adaptor protein Ras-associated and pleckstrin homology domains-containing protein 1, more commonly referred to as Lamellipodin (Lpd). Through its ability to cluster and tether Ena/VASP proteins to actin filaments, Lpd is a key regulator of the actin cytoskeleton^33^. As such Lpd has roles in lamellipodial formation^34^, stabilization of actin-dependent cellular protrusions^35^, cell migration and endocytosis^26, 36–39^. Because the PH domain of Lpd has been shown to bind PtdIns(3,4)P_2_^34^, it appeared a likely candidate for the regulation of pseudopod extension during particle engulfment. To examine this possibility, Lpd was expressed in RAW 264.7 macrophages and its distribution assessed during phagocytosis of IgG-opsonized SRBCs. We observed robust accumulation of GFP-Lpd at the phagocytic cup, where PtdIns(3,4)P_2_ was also enriched (Figure 4A). Furthermore, upon treatment with wortmannin the levels of both PtdIns(3,4)P_2_ and Lpd at the phagocytic cup decreased in parallel fashion (Figure 4B,C). Consistent with the notion that Lpd binds to PtdIns(3,4)P_2_ via its PH domain, we found that a GFP-tagged tandem PH domain of Lpd (Lpd- 2xPH) was recruited to the phagocytic cup in cells expressing the inactive INPP4B(C842A)-CAAX (Figure 4D) yet failed to accumulate in cells expressing INPP4B-CAAX. Furthermore, in resting RAW 264.7 cells, Lpd-2xPH showed modest accumulation at the plasma membrane, as we had seen earlier for the cPHx3 and cPHx2 probes, and this limited recruitment was similarly abolished by INPP4B-CAAX, but not by the catalytically-inactive INPP4B(C842A)-CAAX (Figure S3). From these experiments we conclude that Lpd is recruited early during phagocytosis at least in part by its PH domain-mediated interaction with PtdIns(3,4)P_2_.

**Figure 4.**
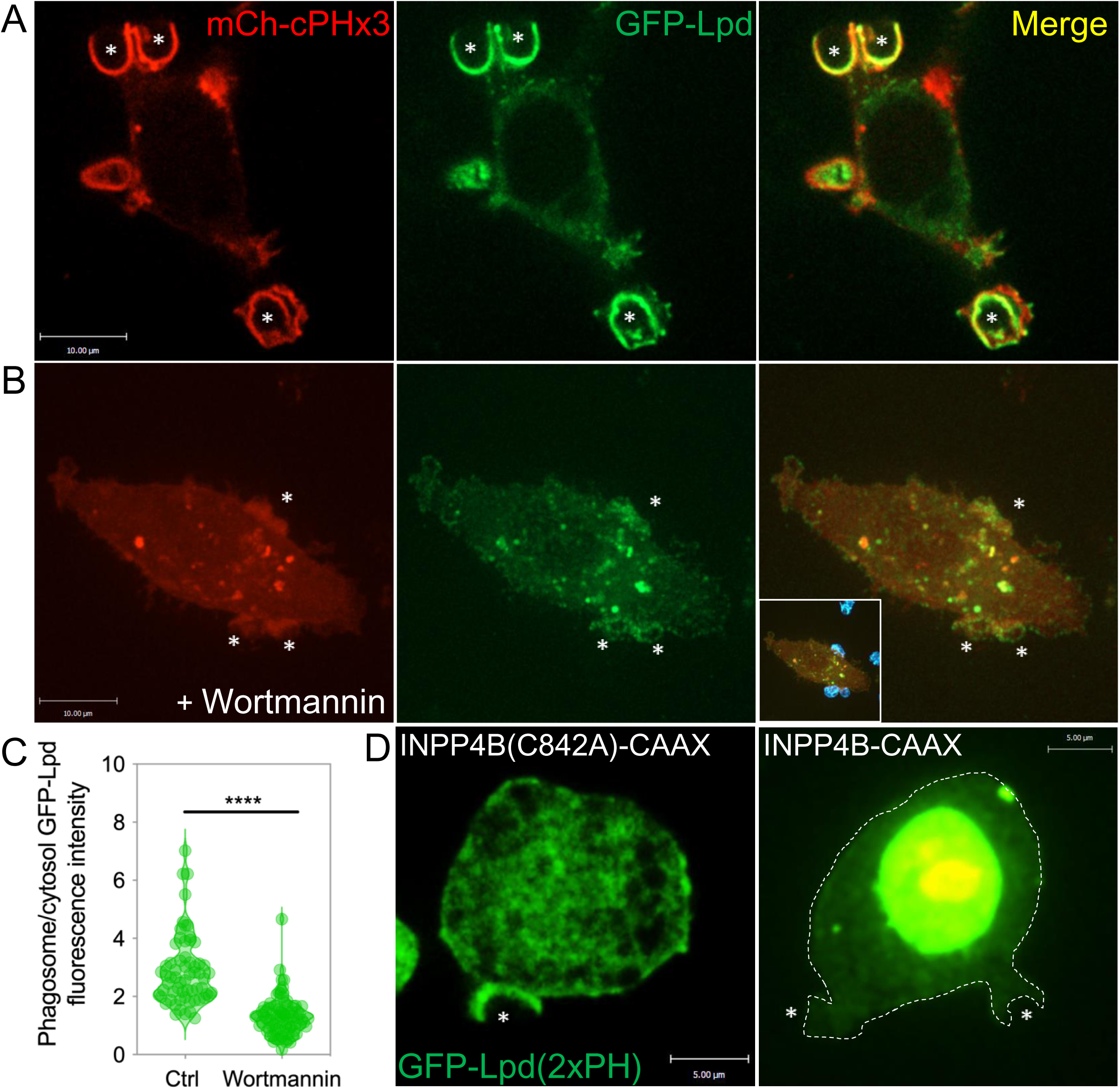
PtdIns(3,4)P_2_ and its effector protein Lamellipodin (Lpd) co-localize at the site of particle engagement during phagocytosis. (A) Representative confocal sections of RAW 264.7 macrophages co-expressing mCherry-cPHx3 and GFP-Lpd during phagocytosis. Asterisks denote sites of phagocytic cup formation. (B) Representative confocal extended-focus projection of RAW 264.7 macrophages co-expressing mCherry-cPHx3 and GFP-Lpd during phagocytosis following pre-treatment with 100 nM wortmannin. Asterisks represent sites of particle engagement. Inset shows engaged SRBCs (C) Normalized phagosomal Lpd fluorescence intensity in control and wortmannin-treated cells. Unpaired *t*-test of individual values across n=3 independent experiments. (D) Representative confocal micrograph of RAW 264.7 cells co-expressing the tandem PH domain of Lpd (green) and either BFP-INPP4B(C842A)-CAAX (left) or BFP-INPP4B-CAAX (right) during phagocytosis. Asterisks represent sites of phagocytosis; dotted line delineates the periphery of the cell. Similar results were observed in 5 independent experiments.

### Silencing of Lpd results in aberrant phagocytic cups and arrests phagocytosis

We next sought to determine whether Lpd is necessary for FcγR-mediated phagocytosis. The expression of endogenous Lpd in RAW 264.7 macrophages and its susceptibility to shRNA-mediated silencing were first validated by immunostaining (Figure 5A and Figure S5). The enrichment of the endogenous Lpd during phagocytosis was similarly confirmed by immunostaining; its distribution at the phagocytic cup closely resembled that of F-actin stained with phalloidin (Figure 5B). Upon silencing of Lpd, we observed an aberrant “flaring” of F-actin around the opsonized particle (Figure 5C, top right panel). Lpd-silenced macrophages exhibited “loose” phagocytic cups characterized by multiple protrusions, in contrast to the tightly apposed pseudopods of control cells. In addition, Lpd-shRNA cells exhibited a ∼60% decrease in phagocytic efficiency compared to control cells (Figure 5D, E). Taken together, these findings suggest that Lpd is necessary for proper F-actin organization and pseudopod extension during phagocytosis.

**Figure 5.**
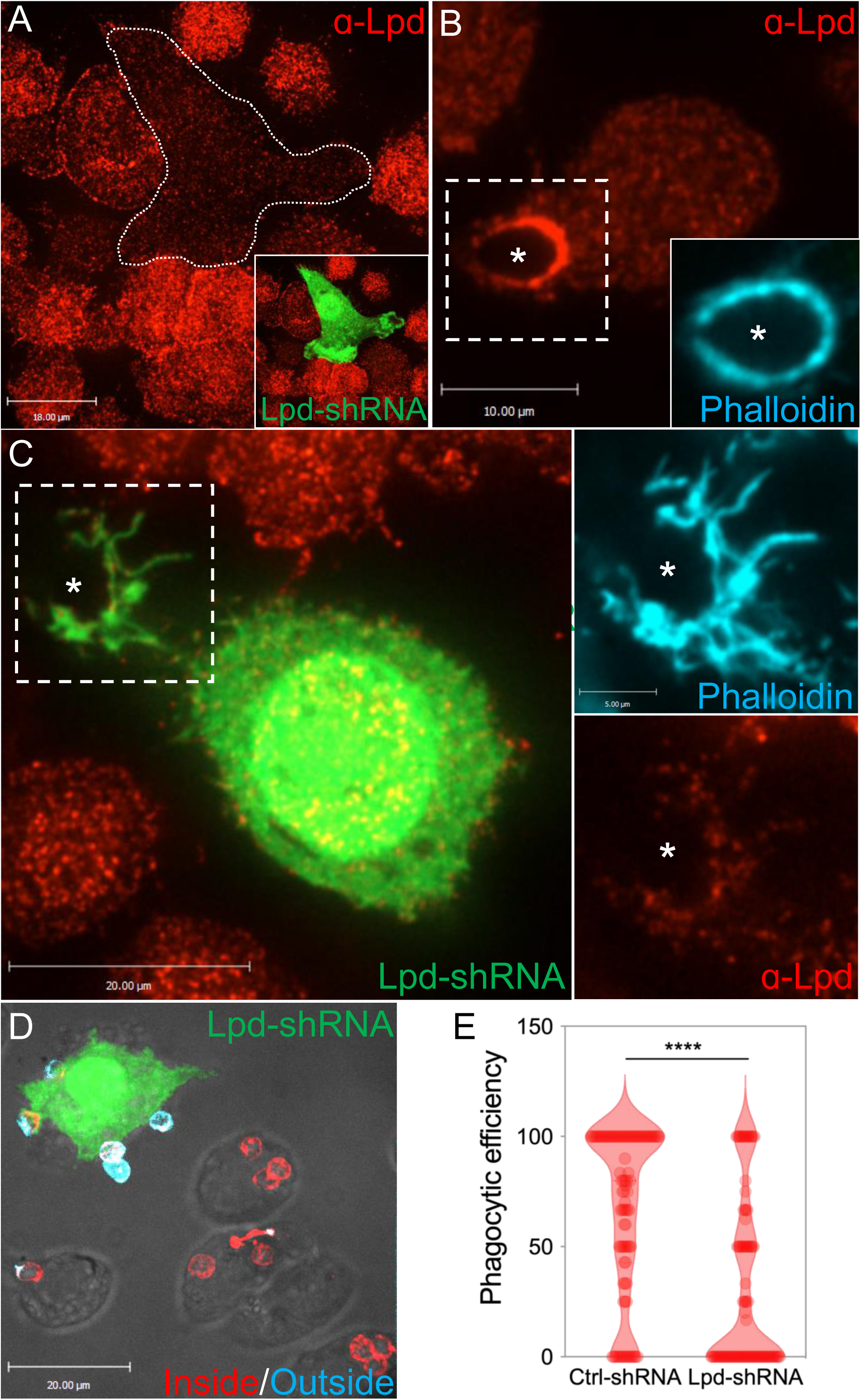
Silencing of lamellipodin in macrophages disrupts phagocytosis and yields an aberrant F-actin morphology at the phagocytic cup. (A) Lpd immunostaining (red) in resting RAW 264.7 macrophages expressing Lpd-shRNA and soluble GFP (green); (B) Lpd immunostaining in control RAW 264.7 macrophages during phagocytosis; (C) Lpd immunostaining in Lpd-silenced RAW 264.7 macrophages (green) during phagocytosis. Asterisks denote sites of particle engagement. Insets are magnifications of the area within the dotted boxes showing Alexa Fluor647-conjugated phalloidin staining (blue) and Lpd staining (red). Similar results were observed in 3 independent experiments. (D) Representative confocal extended-focus projection of RAW 264.7 macrophages with and without expression of Lpd-shRNA (green) following a 10 min incubation with IgG-opsonized SRBCs. Inside-outside staining was performed to differentiate between internalized/total (red) and non-internalized (blue) SRBCs. (E) Quantification of the phagocytic efficiency in Control-shRNA and Lpd-shRNA RAW 264.7 macrophages. Unpaired *t-*test of individual phagocytic efficiency values n=95 (ctrl-shRNA), n=98 (Lpd-shRNA), across N=3 independent experiments; data are means ± SEM.

### The Lamellipodin ligand VASP localizes to the phagocytic cup and is necessary for phagocytosis

Ena/VASP proteins promote actin polymerization by accelerating filament elongation and opposing the action of capping proteins^40–42^. Ena/VASP proteins regulate the cytoskeleton during T cell receptor (TCR)-signalling^43^ and play a role in Fc-mediated phagocytosis^44^ and macroendocytosis in *Dictyostelium discoideum*^45^. It is relevant that Ena/VASP proteins harbour an EVH1 domain that interacts with the multiple proline-rich regions (FPPPP) present in Lpd^33, 34^ (Figure 6A) and that they jointly regulate the dynamics of filopodia^46^. We sought to determine if Lpd and VASP associate within the phagocytic cup. Co-expression of the Lpd and VASP constructs revealed colocalization of these proteins during phagocytosis. Notably, Lpd, VASP and F-actin all share a similar distribution within the phagocytic cup (Figure 6B). We also confirmed the enrichment of endogenous VASP at the phagocytic cup through immunostaining (Figure 6E).

**Figure 6.**
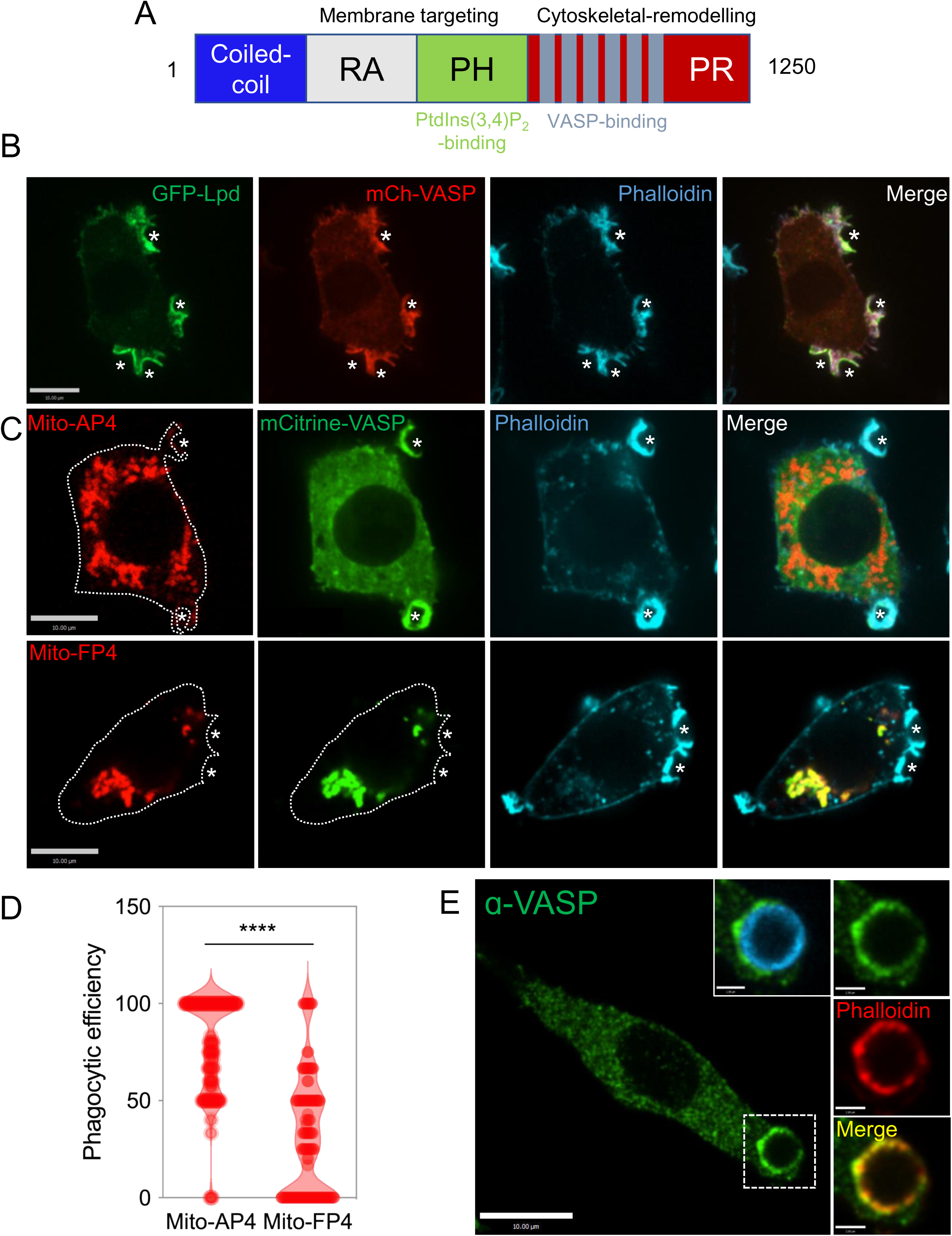
Lamellipodin and VASP co-localize at the phagocytic cup. (A) Schematic representation of the functional domains and binding partners of Lpd. Lpd interacts with PtdIns(3,4)P_2_ and VASP through its PH domain and FP4 repeats respectively. (B) Representative confocal section of RAW 264.7 macrophages co-expressing Lpd (green) and VASP (red) and stained with phalloidin (blue) during phagocytic cup formation. Similar observations were made in 5 independent experiments. Asterisks indicate sites of particle engagement. (C) Representative confocal slice of RAW 264.7 macrophages co-expressing VASP (green) and either the Mito-AP4 (red, top row) or the Mito-FP4 (red, bottom row) constructs. Similar observations were made in three independent experiments. Asterisks indicate sites of particle binding. (D) Quantification of the phagocytic efficiency in RAW 264.7 macrophages expressing Mito-AP4 and Mito-FP4 . Unpaired *t* test of individual phagocytic efficiency values n=117 (Mito-AP4), n=127 (Mito-FP4) across N=3 independent experiments; data are means ± SEM. (E) Representative confocal micrograph of RAW 264.7 macrophages stained for VASP (green) and phalloidin (red) during phagocytosis. Insets represent the area denoted by the dotted box and particle engagement (blue).

Next, we tested whether VASP was necessary for phagocytosis. To this end, we took advantage of the *Listeria monocytogenes* effector protein ActA which binds tightly to VASP. We expressed an N-terminally truncated ActA protein which includes the four repeats of the VASP-binding sequences attached to a C-terminal motif that targets the protein to the cytosolic surface of mitochondria (mRFP-Mito-FP4)^47, 48^. Expression of mRFP-Mito-FP4 effectively sequestered the vast majority of VASP to the mitochondria and prevented its accumulation at the phagocytic cup, while a mutant version (mRFP-Mito-AP4) that also targets to mitochondria but is unable to bind VASP was without effect (Figures 6C and S6). Tethering of VASP to the mitochondria by means of mRFP-Mito-FP4 reduced the phagocytic efficiency from 79.8% to 31.4% (Figure 6D). These results demonstrate that VASP is a positive regulator of phagocytosis and support the notion that Lpd is exerting its effects at least in part by binding to VASP.

### Lamellipodin-VASP interactions coordinate actin polymerization at the phagocytic cup

Next, we examined whether the interaction between Lpd and VASP was necessary for phagocytosis. We overexpressed a mutant version of Lpd in which all Ena/VASP-binding sites had been inactivated through mutations (Lpd^EVmut^). Overexpression of GFP-Lpd^EVmut^ produced a ∼57% decrease in phagocytosis when compared to RAW 264.7 cells transfected with the wild-type GFP-Lpd construct (Figure 7A-B). Furthermore, even though the initial recruitment of GFP- Lpd^EVmut^ to the phagocytic cup was unaffected, the distribution of the construct became altered during phagocytic cup formation (Figure S7). We noticed that the phagocytic cups formed by this mutant contained multiple filopodia- and ruffle-like projections that were rich in F-actin, as visualized through phalloidin staining (Figure 7C). These atypical phagocytic cups and pseudopods closely resembled the ones we had previously observed upon silencing of Lpd (Figure 5C). Additionally, these aberrant phagocytic cups were also enriched in PtdIns(3,4)P_2_, detected by the cPHx3 probe (Figure 7D) demonstrating that the lipid metabolism itself was unperturbed by expression of the Lpd mutant. These findings suggest that interactions between Lpd and VASP are required for proper cytoskeleton organization and coordinated extension of pseudopods at the site of particle engulfment.

**Figure. 7.**
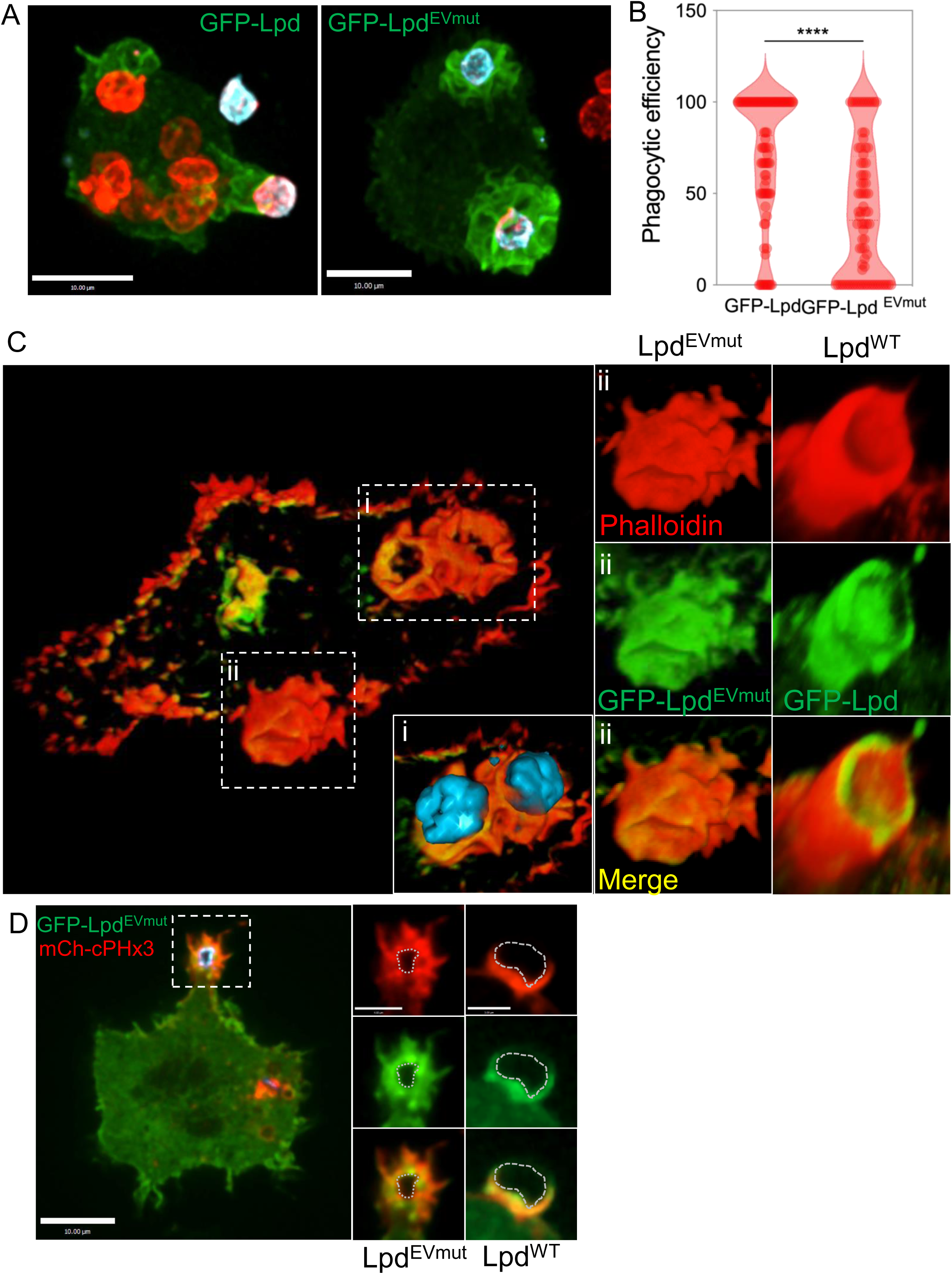
Lamellipodin and VASP coordinate actin polymerization at the phagocytic cup. (A) Representative confocal projections of RAW 264.7 macrophages expressing GFP-Lpd (left) and GFP-Lpd^EVmut^ (right) after a 10-min incubation with IgG-opsonized SRBCs. Inside-outside staining was performed to differentiate between internalized/total (red) and non-internalized (blue) SRBCs. (B) Quantification of the phagocytic efficiency in RAW 264.7 macrophages overexpressing either Lpd^WT^ or Lpd^EVmut^. Unpaired *t* test of individual phagocytic efficiency values n=86 (Lpd^WT^), n=80 (Lpd^EVmut^), across N=3 independent experiments; data are means ± SEM. (C) Representative 3D reconstruction of RAW 264.7 macrophages expressing the Lpd^EVmut^ construct (green) during phagocytosis of IgG-opsonized SRBCs and stained with phalloidin (Red) (i) Magnification of the area denoted by the dotted box (i) in main panel, depicting SRBCs (blue). (ii) Magnification of the area denoted by the dotted box (ii) in main panel, depicting a representative phagocytic cup from RAW 264.7 macrophages expressing either the Lpd^EVmut^ (left) or the Lpd^WT^ construct (right) for comparison. Similar observations were made in 3 independent experiments. (D) Representative confocal slice of RAW 264.7 macrophages co- expressing the Lpd^EVmut^ construct (green) and the cPHx3 construct (red) during phagocytosis. Insets represent the area denoted by the dotted box (left) and compared to the Lpd^WT^- expressing macrophages (right).

## Discussion

We found that comparatively small amounts of PtdIns(3,4)P_2_ are present in the plasma membrane of resting RAW 264.7 macrophages and that this phosphoinositide is greatly enriched at the sites of particle engagement during FcγR-mediated phagosome formation. Selective depletion using a highly specific phosphatase revealed that PtdIns(3,4)P_2_ has a critical role in supporting phagocytosis. This was previously unrecognized in part due to the use of probes like AKT-PH that have dual specificity and of PI3K inhibitors, rather than targeted enzymatic depletion by INPP4B. We also report that Lpd and its binding partner VASP are jointly required for robust phagocytosis: loss of either from the site of phagocytosis results in an atypical and seemingly uncoordinated actin assembly within the extending pseudopods that hence fail to encircle the target.

The importance of PI3Ks during the process of phagocytosis has been widely documented^49,50,51^. Genetic manipulation of PI3Ks as well as the use of pharmacological inhibitors impairs phagocytosis, particularly that of larger targets. Additionally, localized production of PtdIns(3,4)P_2_ or PtdIns(3,4,5)P_3_ at the phagocytic cup has been implicated in phagosomal closure and Ca^2+^ signalling in HL60 neutrophils^10^. Although some of these defects can be directly linked to the role of PtdIns(3,4,5)P_3_ cognate effectors, based on this study it is likely that a second, underappreciated effect of PI3K inhibition is the impairment of PtdIns(3,4)P_2_ formation, which is itself required for phagocytosis. Curiously, in our studies we find that PtdIns(3,4)P_2_ is found at both the base of the phagocytic cup as well as at the tips of the pseudopods, whereas PtdIns(3,4,5)P_3_ is largely restricted to the base of the phagocytic cup (Figure S1). This raises the possibility that the relevant phosphoinositide 5-phosphatases are particularly active at the tips of the advancing pseudopods. Phenotypically, inhibition of PI3K – and thus attenuation of both PtdIns(3,4)P2 and PtdIns(3,4,5)P3 production– results in arrest of pseudopod progression roughly halfway around the opsonized particle^9, 52^, while selective depletion of PtdIns(3,4)P_2_ prevents pseudopod extension (Figure 3C right panel, 3D, and Video S3). PtdIns(3,4,5)P_3_ is thought to regulate actin dynamics during phagocytosis by locally activating Rho family GEFs and GAPs^53^, and by activating phospholipase Cγ, thereby depleting PtdIns(4,5)P_2_ from the base of the phagocytic cup. In view of our observations, it seems worthwhile to reconsider whether the purported PtdIns(3,4,5)P_3_ effectors are in fact selective for this lipid or are instead responsive to PtdIns(3,4)P_2_.

An earlier report suggested that the inositol polyphosphate 4-phosphatase type 1 (INPP4A), which is expressed in RAW macrophages, is a negative regulator of phagocytosis^54^. That study documented enhanced ability to internalize particles by RAW cells expressing shINPP4A and by primary macrophages from *INPP4A*-knockout mice. While the authors did not provide a mechanistic explanation for the observed effects, their findings are consistent with our conclusion that increased PtdIns(3,4)P_2_ is essential for optimal phagocytosis, at least partly via Lpd and VASP.

Lpd belongs to the <u>M</u>ig-10, <u>R</u>IAM, <u>L</u>pd (MRL) family of modular adaptor proteins^55^. RIAM, despite sharing homology with Lpd^56^, is thought to act downstream of Rap1, and evidence suggests that its PH domain binds preferentially to PtdIns(3)P, PtdIns(5)P ^57^ and PtdIns(4,5)P_2_^58, 59^, not PtdIns(3,4)P_2_. Previously, RIAM was described to support complement-dependent phagocytosis by relaying integrin signaling^60^. However, RIAM is not required for FcγR-dependent phagocytosis^61^, the mode we studied. Furthermore, our results indicate that silencing of Lpd produces severe defects in the phagocytic cups of macrophages where RIAM was left intact. These findings suggest that Lpd, contrary to RIAM, is necessary for Fc-mediated phagocytosis and that these MRL proteins play non-redundant roles during this process.

We found that the PH domain of Lpd is sufficient for its localization at the phagocytic cup. It is possible however, that other interactions contribute to its recruitment and retention within the forming phagosome. Indeed, we noted that Lpd dissociates from the maturing phagosome prior to the disappearance of the PtdIns(3,4)P_2_, suggesting that it may function as a co- incidence detector; other potential binding partners may well be present at the site of phagocytosis. In this regard, it is noteworthy that Lpd directly binds actin filaments through an unstructured and highly basic region in its carboxyl-terminus^33^. Furthermore, Lpd can also bind to active Rac1^62, 63^, which is similarly recruited to forming phagocytic cups^64^. Additionally, the formin-binding protein 17 (FBP17) binds to Lpd and recruits it to the membrane during fast endophilin-mediated endocytosis^65^. Of note, FPB17 is also present within podosomes and phagocytic cups in macrophages^66^.

Lpd is central to the regulation of cytoskeletal dynamics^34^ as it interacts with multiple actin regulators such as the Ena/VASP family proteins^33, 37, 67^ and the SCAR/WAVE regulatory complex^37, 62^. Ena/VASP proteins localize to the phagocytic cup and are indispensable for phagocytosis: Coppolino *et al*.^44^ reported that upon binding of Ena/VASP to a cytosolic GFP-ActA construct, phagocytosis was inhibited. In the current study, we were able to validate these findings using a more thorough method that entailed mistargeting Ena/VASP proteins to the mitochondria. Additionally, we found that all three Ena/VASP family members (EVL, Mena, VASP) localize to the phagocytic cup when ectopically expressed (Figure S8). Phosphorylation of Lpd by c-Abl kinase reportedly increases its interaction with Ena/VASP ^68^. While we did not explore this possibility in the current study, Abl family kinases have been implicated in the positive regulation of phagocytosis^69^. Therefore, it is possible that a kinase-dependent increase in Lpd–Ena/VASP interaction is one of the mechanisms by which this positive regulation takes place. Furthermore, PI3K activation through Abl kinase was also recently reported to take place during podosome formation in macrophages^70^.

In addition to VASP, Lpd can also regulate the actin cytoskeleton through its interactions with the SCAR/WAVE complex. A recent study demonstrated that loss of the WAVE regulatory complex (WRC) impairs phagocytosis^71^. In our study, we compared the effects of Lpd mutants in which all Ena/VASP or all SCAR/WAVE binding sites had been mutated. The Lpd^EVmut^ caused the most robust decrease in phagocytic efficiency and aberrant phagocytic cups (Figure 7).

Nevertheless, the SCAR/WAVE binding-deficient mutant also exhibited a smaller, yet significant, decrease in phagocytic efficiency (Figure S9B and C). Furthermore, Abi1, the component of the WRC which interacts directly with Lpd ^62^, was also localized to the phagocytic cup (Figure S9A). Jointly, these findings suggest that Lpd is central to the regulation of actin dynamics during phagocytic cup formation not only through its interactions with VASP, but probably also through the regulation of the WRC, although additional experiments will be needed to validate the involvement of the latter pathway.

The diversity of phagocytic targets and receptors entails different molecular mechanisms during phagocytic cup formation. Despite these differences, all types of phagocytosis share an inherent dependence on the rearrangement of the actin cytoskeleton and on the dynamic remodelling of the plasma membrane. FcγR-mediated phagocytosis is the best characterized model of phagocytosis and the one used throughout this study. However, other equally important modes of opsonin-dependent and independent phagocytosis exist. These include phagocytosis mediated by CR3, receptors that recognize activated complement components, such as iC3b, deposited on the phagocytic target^72^. As in the case of FcγR-mediated phagocytosis, we were able to document that localized synthesis of PtdIns(3,4)P_2_ and recruitment of Lpd and VASP occur also at phagocytic cups generated by engagement of CR3 (Figure S10A, B and C). Interestingly, the overall efficiency of complement-induced phagocytosis was largely unaffected by expression of INPP4B (Figure S10D). One possibility for this difference is compensation by the Lpd homolog RIAM, which is associated with the activation of integrins like CR3 but not FcγR and has the ability to bind multiple phosphoinositides including PtdIns(3,4,5)P_3_ and PtdIns(4,5)P_2_ ^59^, and recently shown to be required for CR3-mediated phagocytosis in HL60 cells ^60^. In yet another mode of phagocytosis, the scaffolding protein SWAP70 was identified as a PtdIns(3,4)P_2_ effector critical for the phagocytosis of yeast particles in dendritic cells^73, 74^. Collectively, these results demonstrate that PtdIns(3,4)P_2_ regulates actin dynamics through multiple effectors in at least a subset of phagocytic receptors and cell types.

## Methods

### Plasmids

The plasmids used in this study are summarized in Table 1. Where indicated, plasmids were constructed using In-Fusion HD EcoDry Cloning Kits (Takara) and were verified by Sanger sequencing prior to transfection into mammalian cells.

**Table 1.**

| Plasmid | Backbone vector | Reference |
| --- | --- | --- |
| NES-EGFP-cPHx2 | pNES-EGFP-C1 | 22 |
| NES-EGFP-cPHx3 | pNES-EGFP-C1 | 22 |
| NES-mCherry-cPHx3 | pNES-mCherry-C1 | 22 |
| NES-mCherry-PH-BTKx2 | pNES-mCherry-C1 | Walpole et al., Nature Cell Biology 2022 |
| EGFP-P4Mx2 | pEGFP-C1 | 77 |
| mCherry-P4Mx2 | mCherry-C1 | 77 |
| TagBFP2-INPP4B<sup>WT</sup>-CAAX | pTagBFP2-C1 | 22 |
| TagBFP2-INPP4B<sup>C842A</sup>-CAAX | pTagBFP2-C1 | 22 |
| pSBtet-pur-TagBFP2-INPP4B(WT)-CAAX | pTagBFP2-C1 | This study |
| pSBtet-pur-TagBFP2-INPP4B(C842A)-CAAX | pTagBFP2-C1 | This study |
| pSBtet-pur |  | PMID: 25650551; Addgene #60507 |
| pSBtet-BP |  | PMID: 25650551; Addgene #60496 |
| pCMV(CAT)-T7-SB100 |  | PMID: 19412179; Addgene #34879 |
| NES-mCherry-PH-BTKx2 |  | This study |
| GFP-Lpd<sup>EVmut</sup> | pEGFP-C1 | 62 |
| GFP-LpdS/Wmut | pEGFP-C1 | 62 |
| GFP-Lpd | pEGFP-C1 | 34 |
| Lifeact-EGFP | pEGFP-C1 | PMID: 26317264; Addgene #58470 |
| GFP-Lpd(2xPH) | pEGFP-C1 | 34 |
| GFP-Lpd(ΔPH) | pEGFP-C1 | 78 |
| mRFP-Mito-FP4 | mRFP | 79 |
| mRFP-Mito-AP4 | mRFP | 79 |
| pmScarlet-EVL | pmScarlett | Gift from M. Krause (King's College London) |
| pmScarlet-VASP | pmScarlett | Gift from M. Krause (King's College London) |
| pmScarlet-Mena | pmScarlett | Gift from M. Krause (King's College London) |
| mCitrine-VASP-5 | mCitrine | Addgene #56571 |
| mCherry-VASP-5 | mCherry | Addgene #55151 |
| shRNA mouse Lpd | pLL3.7-Puro | 34 |
| pEGFP-C2-SHIP2 D607A | pEGFPC2 | MRCPPU Reagents, University of Dundee |
| GFP-SHIP2 | pEGFPC2 | MRCPPU Reagents, University of Dundee |
| Inpp5B-GFP | pEGFP-C1 | 80 |
| pEGFP-Abi1 | pEGFP-C1 | Addgene #74905 |

### Cell Culture

The murine macrophage RAW 264.7 cell line was obtained from the American Type Culture Collection (Manassas, VA) and cultured in DMEM (Wisent Bioproducts, Saint-Jean-Baptiste, Canada) supplemented with 10% heat-inactivated fetal bovine serum and incubated at 37°C under 5% CO_2_.

### Phagocytosis assays

Quantitative phagocytosis assays have been described previously^75^. Briefly, ∼5 × 10^4^ cells were seeded onto 1.8-cm glass coverslips and grown for 18–24 h. Opsonization of SRBC was performed by incubating 100 μL of a 10% SRBC suspension with 3 μL of rabbit IgG fraction against SRBCs at 37°C for 1 h. SRBCs were then labelled with a fluorescent secondary antibody against rabbit. 10 μL of labelled and opsonized SRBCs was added to individual coverslips containing RAW cells within 12-well plates or directly into imaging chambers for live-cell imaging. Synchronization of phagocytosis was achieved by centrifugation (300 ×*g* for 10 s) of the phagocytic targets onto the cell-containing coverslips. Phagocytosis was terminated by replacing the medium with ice-cold PBS. To obtain the differential inside-outside staining used throughout this study, cells were incubated with cold PBS containing a secondary antibody against rabbit to label non-internalized SRBCs, previously internalized labelled-SRBCs are not accessible to this secondary antibody staining and therefore are not additionally labelled.

### Antibodies and reagents

Rabbit polyclonal Lpd antibody^34^ was used for immunofluorescence at 1:200. Rabbit monoclonal VASP (9A2) antibody (Cell Signaling Technology, #3132) was used for immunofluorescence at 1:200. SRBCs 10% suspension was purchased from MP Biomedicals (Santa Ana, CA). Anti-sheep red blood cell antibodies were purchased from Cedarlane Laboratories (Burlington, Canada). Fluorescent secondary antibodies against mouse and rabbit were purchased from Jackson ImmunoResearch Labs (West Grove, PA). Paraformaldehyde (16% w/v) was purchased from Electron Microscopy Sciences (Hatfield, PA). Wortmannin (Sigma 681675), PI-103 (Sigma 528100), GDC-0941 (Millipore Sigma 509226), GSK-A1 (kindly provided by Dr. Tamas Balla^25^), AS1949490 and K118 (Echelon Biosciences). Fluorescent phalloidin (Molecular Probes). Phosphate-buffered saline (PBS) and Hanks’ balanced salt solution (HBSS) (Wisent Bioproducts, Saint-Jean-Baptiste, Canada.

### Transfections

RAW 264.7 cells were plated in 18 mm round coverslips ∼36–40 h before experiments. Subsequently, cells were transfected using Fugene HD (Promega) according to the manufacturer’s protocol. Briefly, 1 μg of plasmid DNA was mixed with 3 μL of Fugene HD in 100 μL of serum-free DMEM. The mix was incubated for 15 min at room temperature, then distributed into two wells of a 12-well plate containing cells seeded onto glass coverslips. Experiments were performed ∼16–18 h post-transfection.

### Inducible cell line generation

Transgenic, doxycycline-inducible RAW 264.7 stable lines were generated by the Sleeping Beauty transposon system^76^. The open reading frames of TagBFP2-INPP4B(WT)-CAAX or TagBFP2-INPP4B(C842A)-CAAX were inserted into the pSBTet-pur backbone. pSBTet-pur plasmids were co-transfected in a 10:1 ratio to pCMV(CAT)-T7-SB100 in 6-well plates using FuGENE HD as described above. Cell culture medium was replaced 18 hours later and, to select for genomic integration, RAW 264.7 cells were incubated with 2.5 μg/mL puromycin (Sigma P8833). Puromycin selection was maintained for 5 days prior to expanding the polyclonal population into culture flasks for expansion and validation. For induction of TagBFP2-fusions, 1 μg/mL doxycycline (Hyclate, Sigma D9891) was added to the culture medium for 48 hours.

Generally, ∼50% of the polyclonal RAW 264.7 cell population exhibited robust expression of the transgenes.

### Immunostaining

Paraformaldehyde (4% w/v)-fixed cells were permeabilized in 0.1 % (v/v) Triton X-100 in PBS for 7 min, blocked in 2% (w/v) BSA in PBS for 1 hour at room temperature or overnight at 4°C. Then, coverslips were incubated for 1 hour with primary antibodies (diluted in 2% BSA), washed multiple times in PBS and subsequently incubated with secondary antibodies (diluted in PBS) for 1 hour.

### Confocal microscopy and image processing

Confocal fluorescence microscopy was performed using spinning-disk confocal microscopes (Quorum Technologies). Our systems use an Axiovert 200M microscope (Carl Zeiss) equipped with a ×63 oil immersion objective (NA 1.4) and a ×1.5 magnifying lens. These microscopes carry a motorized XY stage (Applied Scientific Instrumentation), a piezo Z-focus drive and diode-pumped solid-state lasers emitting at 440, 491, 561, 638 and 655 nm (Spectral Applied Research). Images were recorded with back-thinned, cooled charge-coupled device cameras (Hamamatsu Photonics). Acquisition settings and capture are controlled by Volocity software v6.2.1 (PerkinElmer). Selection of regions of interest, fluorescence intensity measurements and brightness–contrast corrections were performed with Volocity software or with FIJI (National Institutes of Health). Brightness and contrast parameters were adjusted across entire images without altering the linearity of mapped pixel values. For live-cell imaging, cells grown on glass coverslips were mounted in a Chamlide magnetic chamber (Seoul, South Korea), filled with 1 mL of HBSS and kept at 37°C with a heating stage.

### Statistics and reproducibility

Statistical analyses were performed using GraphPad PRISM. Statistical details for each experiment are also stated in the figure legends. Statistical significance (****P < 0.0001, ***P < 0.001, **P < 0.01, *P < 0.05 and NS (not significant)) is denoted in figures when applicable. All microscopy-based experiments were performed independently at least three times.

## Supplemental Information

**Supplemental Figure 1.**
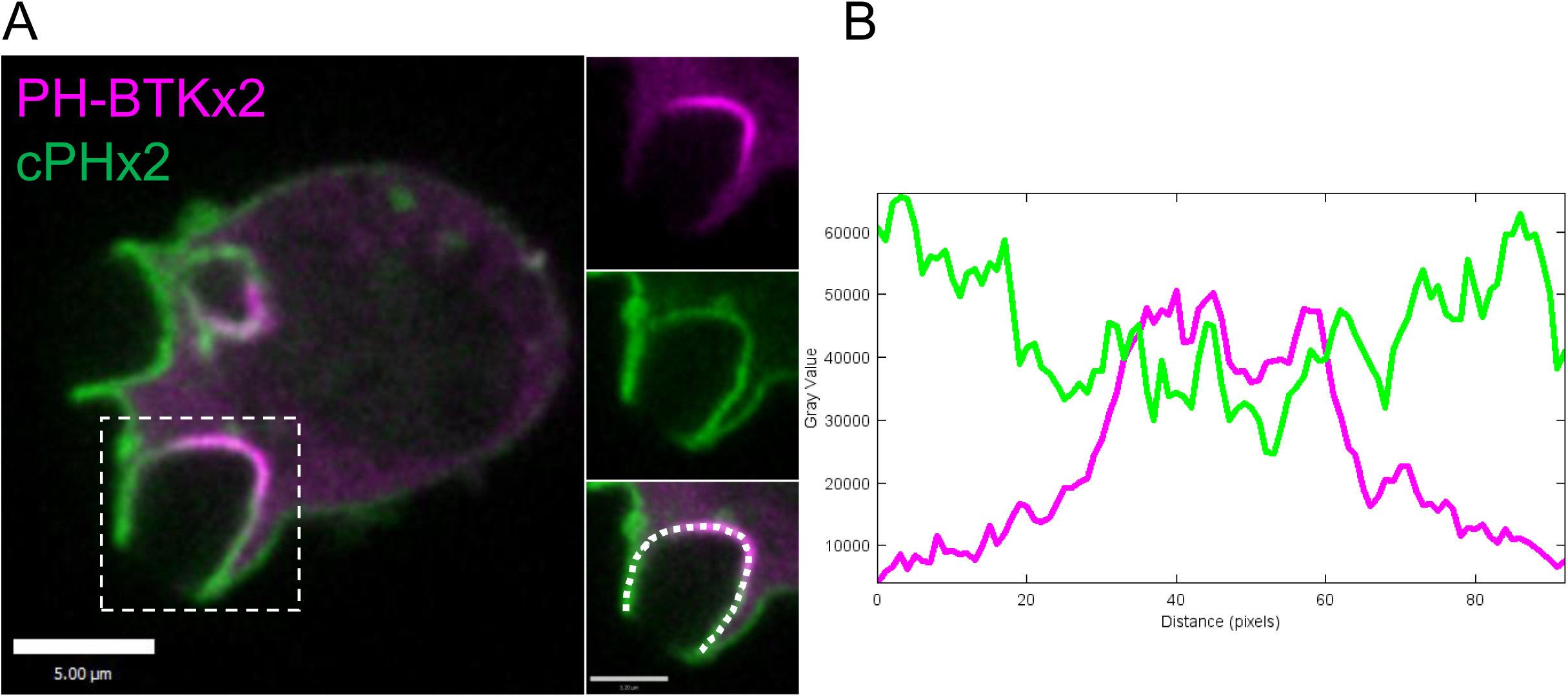
PtdIns(3,4)P_2_ and PtdIns(3,4,5)P_3_ are differentially distributed within the phagocytic cup. (A) RAW 264.7 macrophages co-expressing the cPHx2 (green) and PH-BTKx2 (magenta) constructs during phagocytic cup formation. (B) Intensity profile from the dotted line in A of the cPHx2 (green) and PH-BTKx2 (magenta) constructs at the phagocytic cup.

**Supplemental Figure 2.**
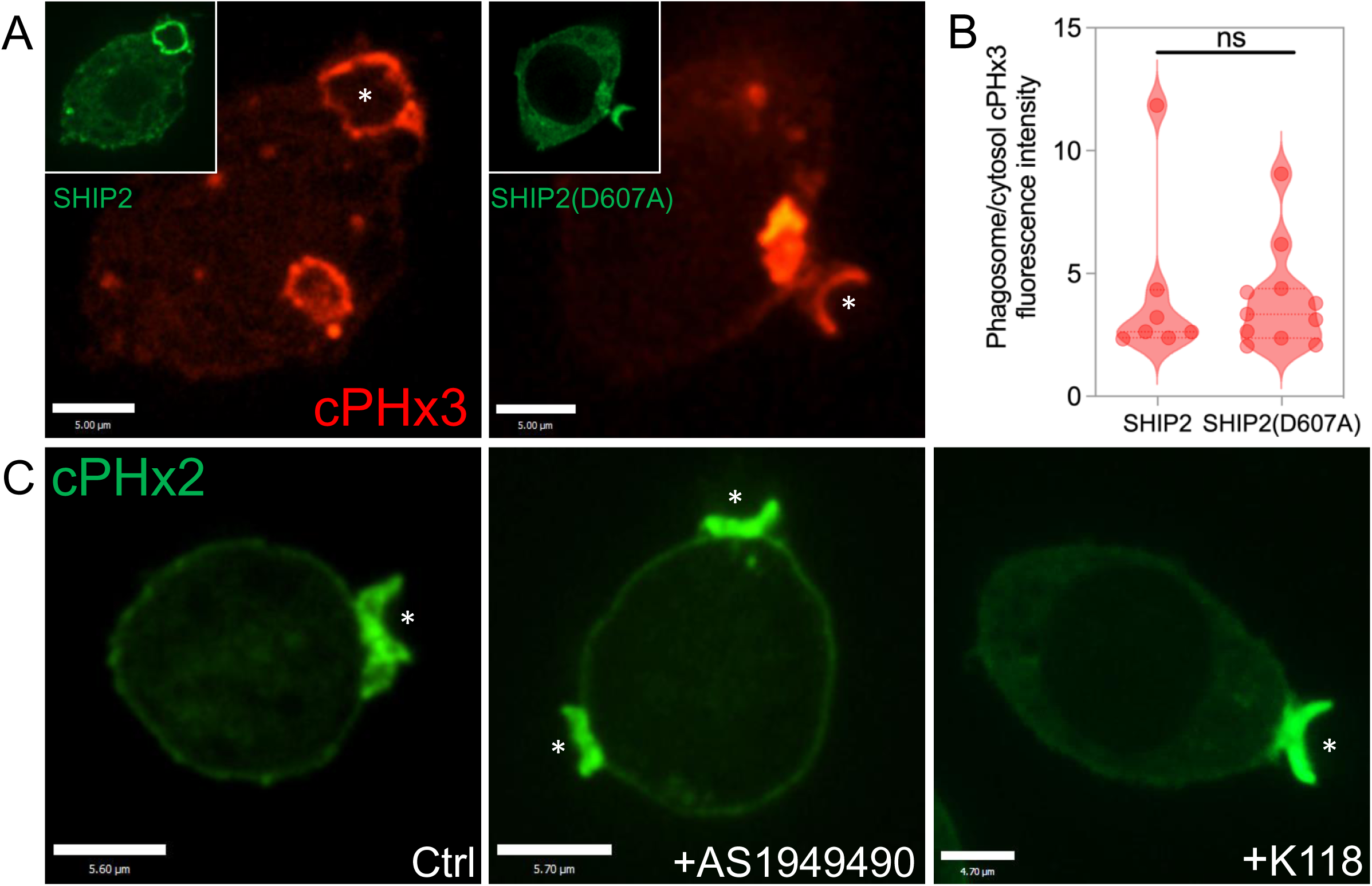
SHIP1 and SHIP2 are dispensable for PtdIns(3,4)P_2_ production at the phagocytic cup. (A) Representative confocal sections of RAW 264.7 macrophages co-expressing the cPHx3 probe (red) with either the wild-type GFP-SHIP2 (left) or the catalytically-inactive GFP- SHIP2(D607A) (right). (B) Normalized phagosomal-cPHx3 fluorescence intensity in cells co-expressing either wild-type GFP-SHIP2 or the catalytically-inactive GFP-SHIP2(D607A). Unpaired *t* test of individual values across n=3 independent experiments; data are means ± SD. (C) Representative confocal slices of RAW 264.7 macrophages expressing the cPHx2 probe (green) in either control cells (left), or cells pre-treated with the SHIP2 inhibitor AS1949490 10 μM for 5h (middle) or pre-treated with the pan-SHIP1/2 inhibitor K118 5 μM for 5h (right). Asterisks indicate sites of phagocytic cup formation.

**Supplemental Figure 3.**
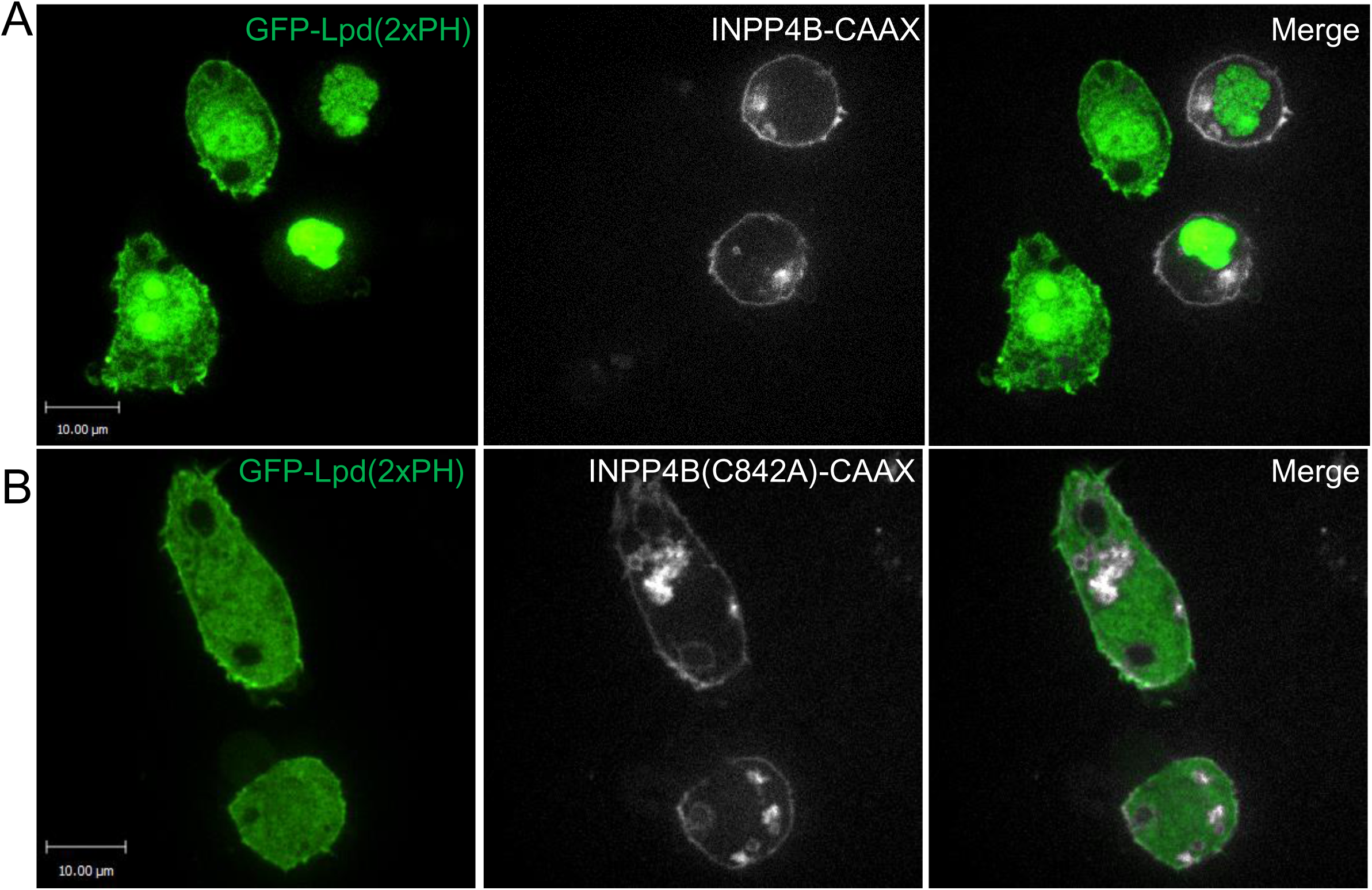
PtdIns(3,4)P_2_ determines the cellular localization of the PH domain of Lpd. (A) Representative confocal sections of RAW 264.7 macrophages expressing the tandem PH domain of Lpd (green) either alone or in combination with the INPP4B-CAAX construct. Similar results were observed in 3 independent experiments. (B) Representative confocal sections of RAW 264.7 macrophages co-expressing the tandem PH domain of Lpd (green) together with the INPP4B(C842A)-CAAX construct. Similar results were observed in 3 independent experimental repeats.

**Supplemental Figure 4.**
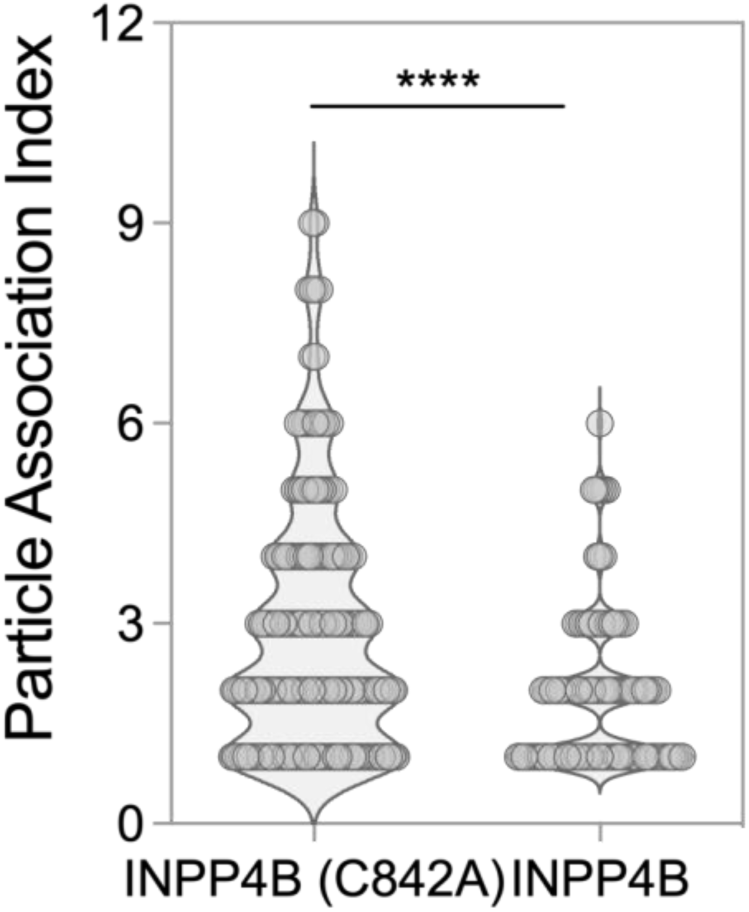
Particle association index in PtdIns(3,4)P_2_ – depleted cells. Quantification of the particle association index in in BFP-INPP4B(C842A)-CAAX and BFP-INPP4B- CAAX-expressing RAW 264.7 macrophages. Unpaired *t*-test of individual particle association index values across N=3 independent experiments; data are mean ± SEM.

**Supplemental Figure 5.**
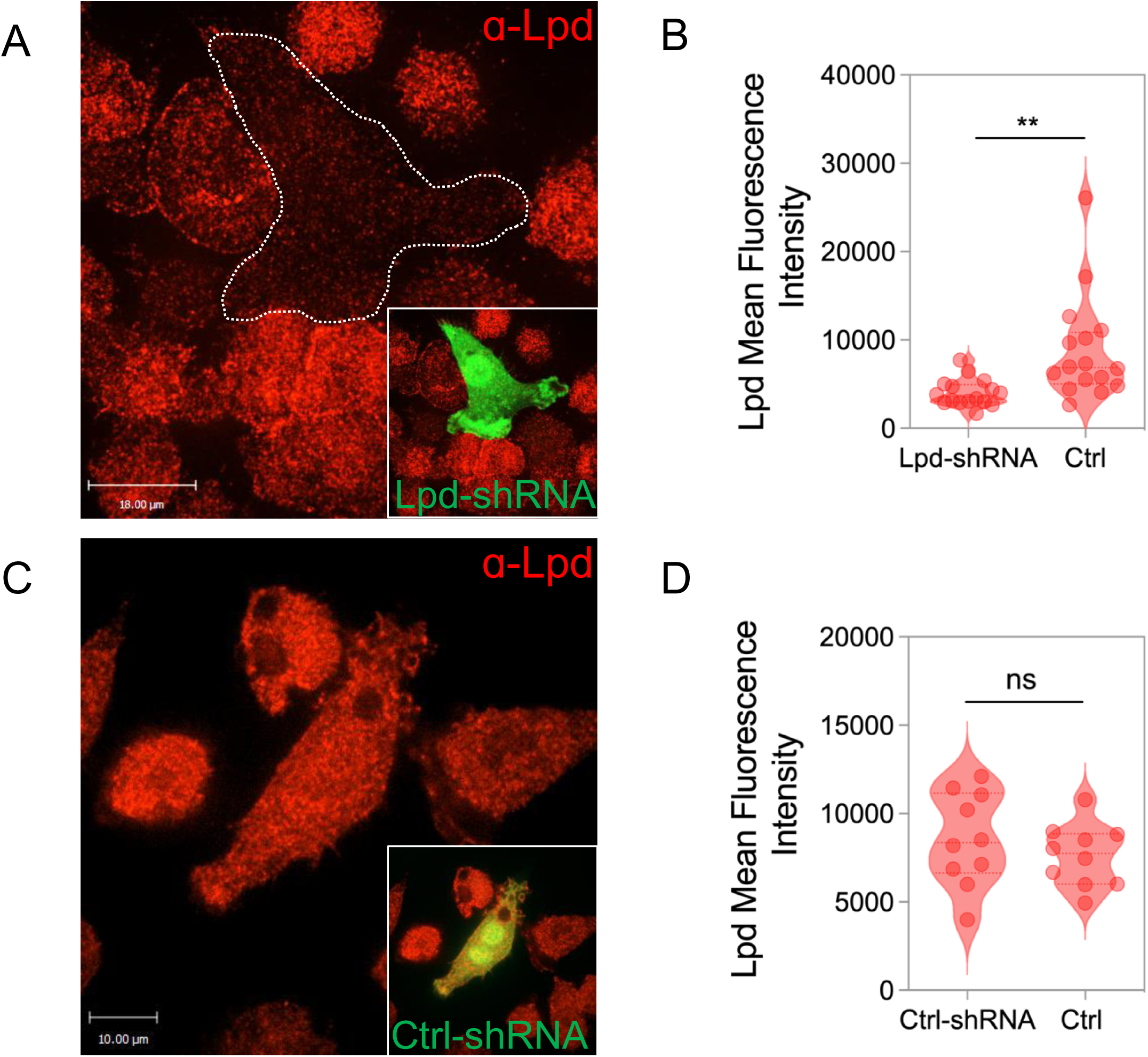
shRNA-mediated silencing of Lamellipodin. (A) Representative confocal micrograph of RAW 264.7 macrophages stained for Lpd (red) and transfected with Lpd-shRNA (green). (B) Quantification of the mean fluorescence intensity of Lpd staining in Lpd-shRNA and control cells. Unpaired *t* test of individual values across n=3 independent experiments; data are means ± SEM. (C) Representative confocal micrograph of RAW 264.7 macrophages stained for Lpd (red) and transfected with Ctrl-shRNA (green). (D) Quantification of the mean fluorescence intensity of Lpd staining in Ctrl-shRNA and control cells. Unpaired *t* test of individual values across n=3 independent experiments; data are means ± SEM.

**Supplemental Figure 6.**
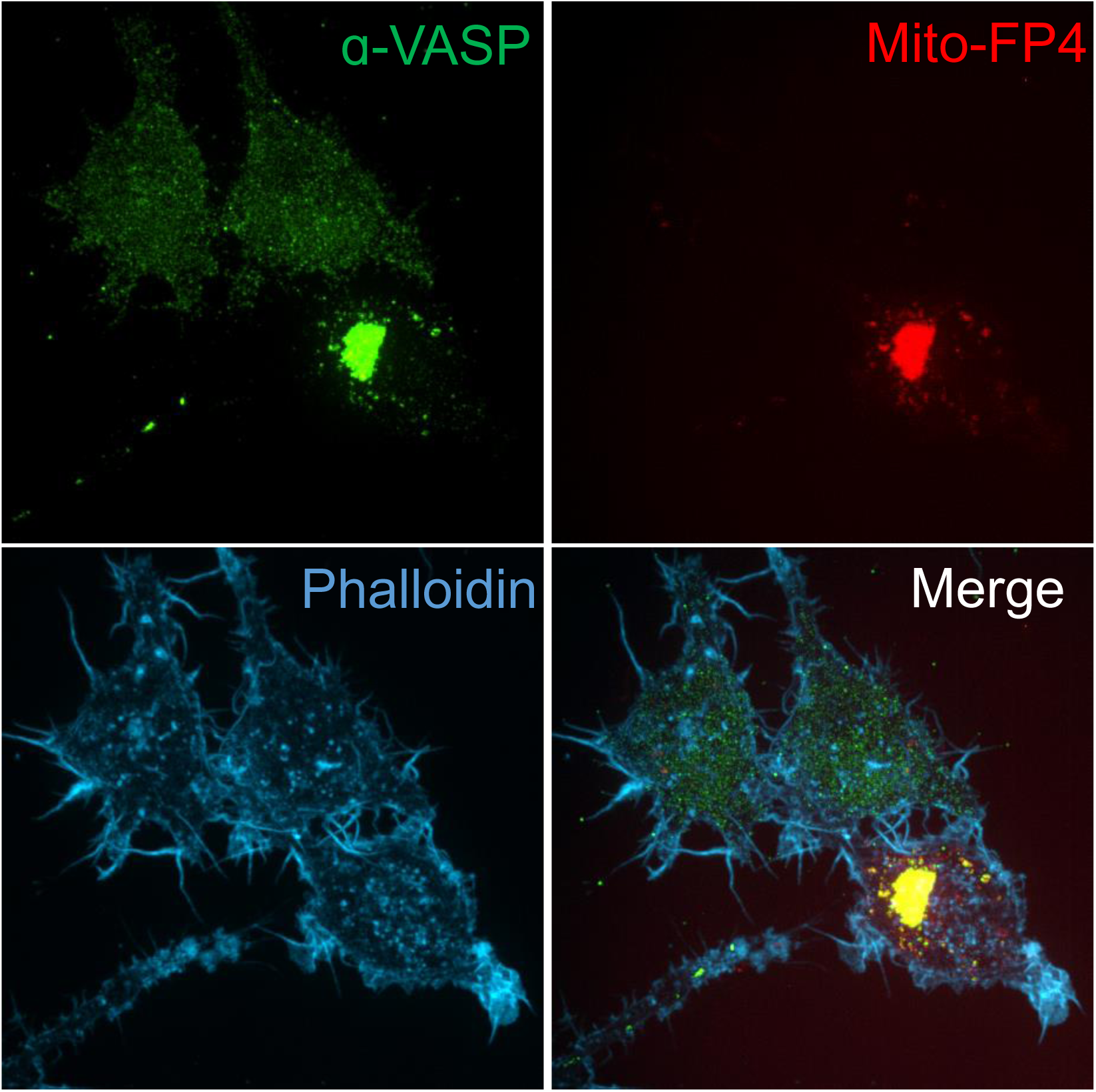
MITO-FP4 targets endogenous VASP to the Mitochondria. Representative confocal extended-focus projections of Raw 264.7 macrophages stained for VASP (green), transfected with the Mito-FP4 construct (red) and stained with phalloidin (blue).

**Supplemental Figure 7.**
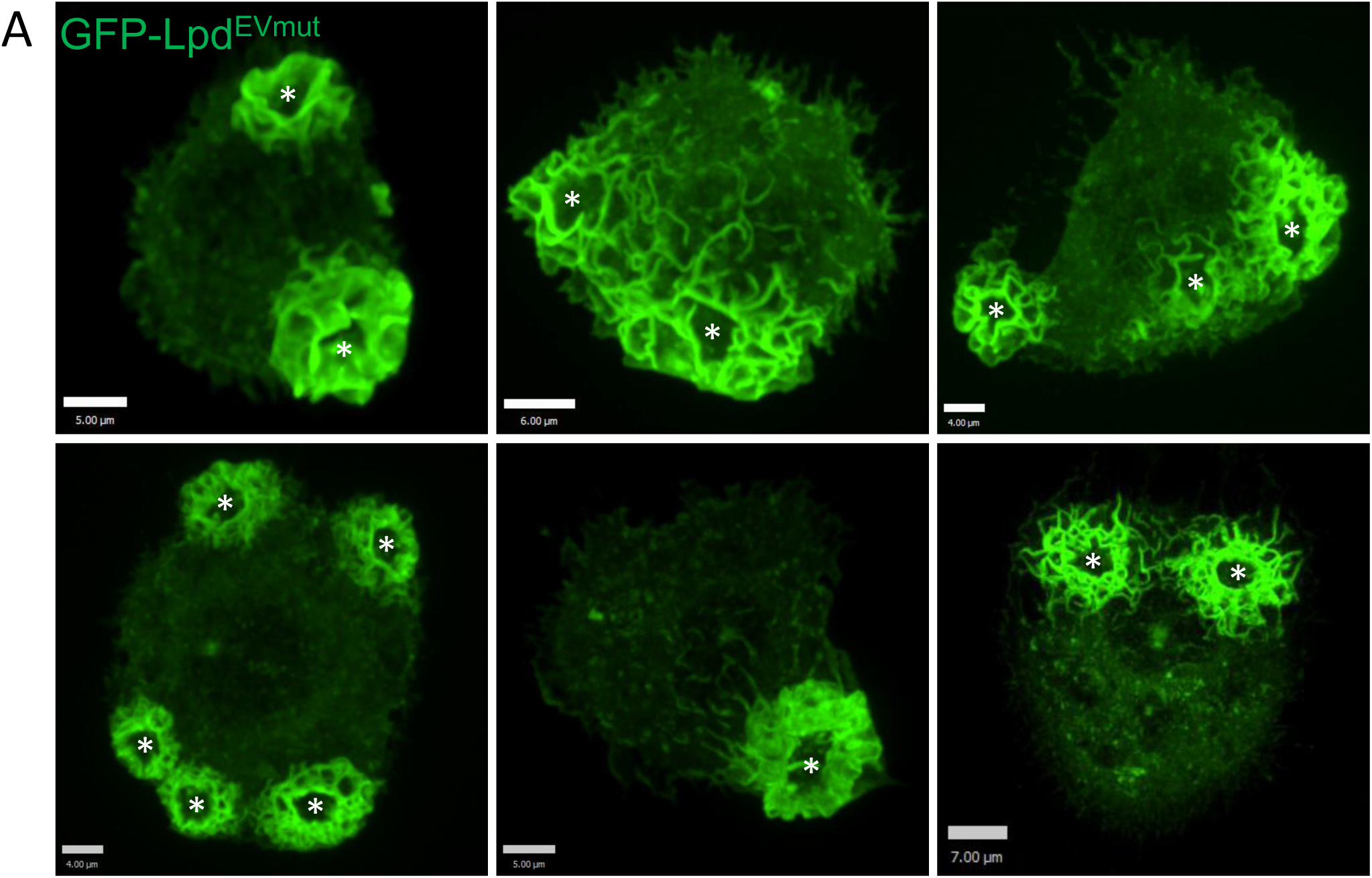
LpdEVmut during phagocytosis. (A) Gallery of representative confocal projections of RAW 264.7 expressing the Lpd^EVmut^ construct during phagocytosis. Asterisks denote sites of phagocytic cup formation.

**Supplemental Figure 8.**
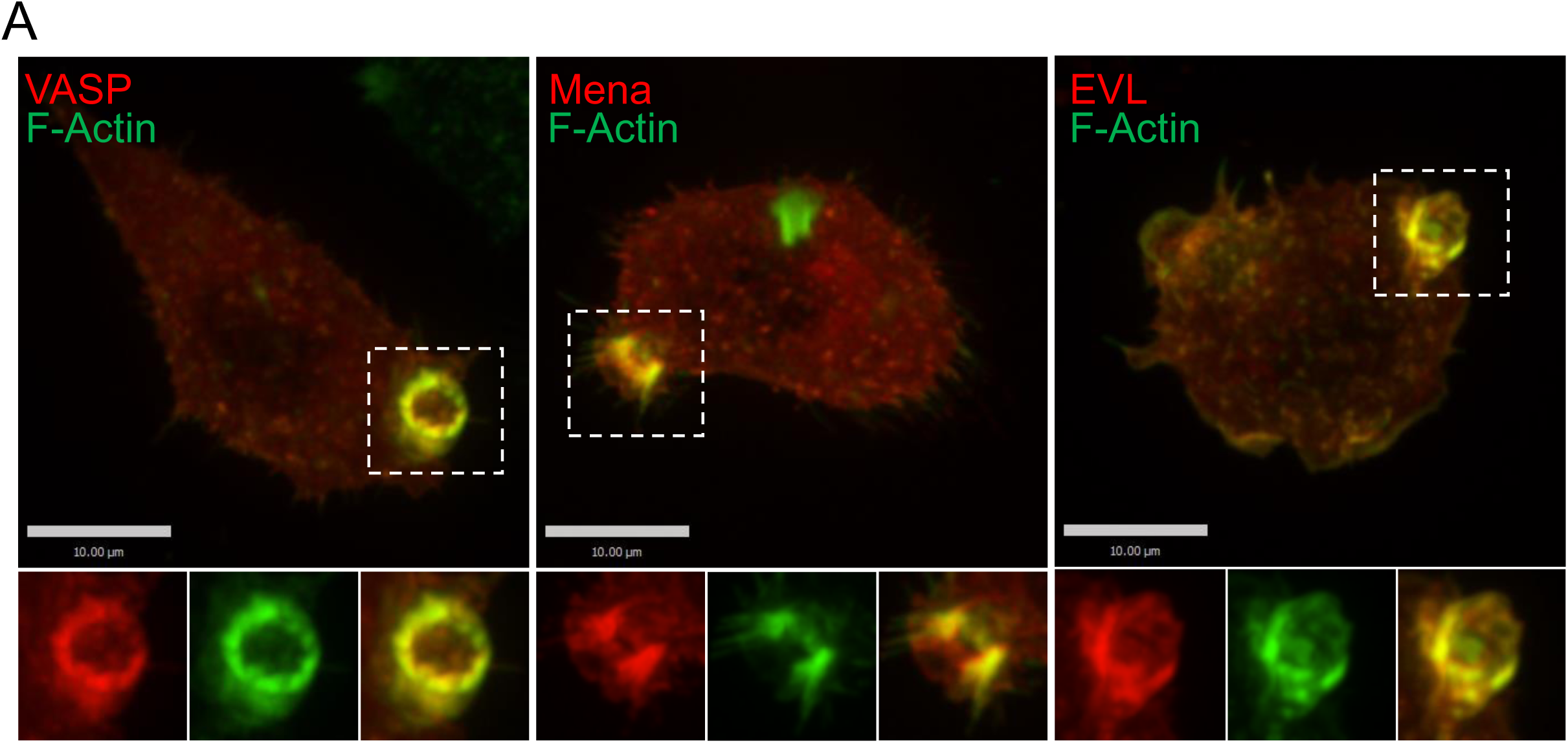
EVL, Mena and VASP localize at the phagocytic cup. (A) Representative confocal sections of RAW 264.7 macrophages stained with phalloidin (green) and expressing VASP (Left), Mena (Middle), and EVL (Right). Insets represent the phagocytic cups denoted by the dotted box. Similar observations were made in 3 independent experimental repeats.

**Supplemental Figure 9.**
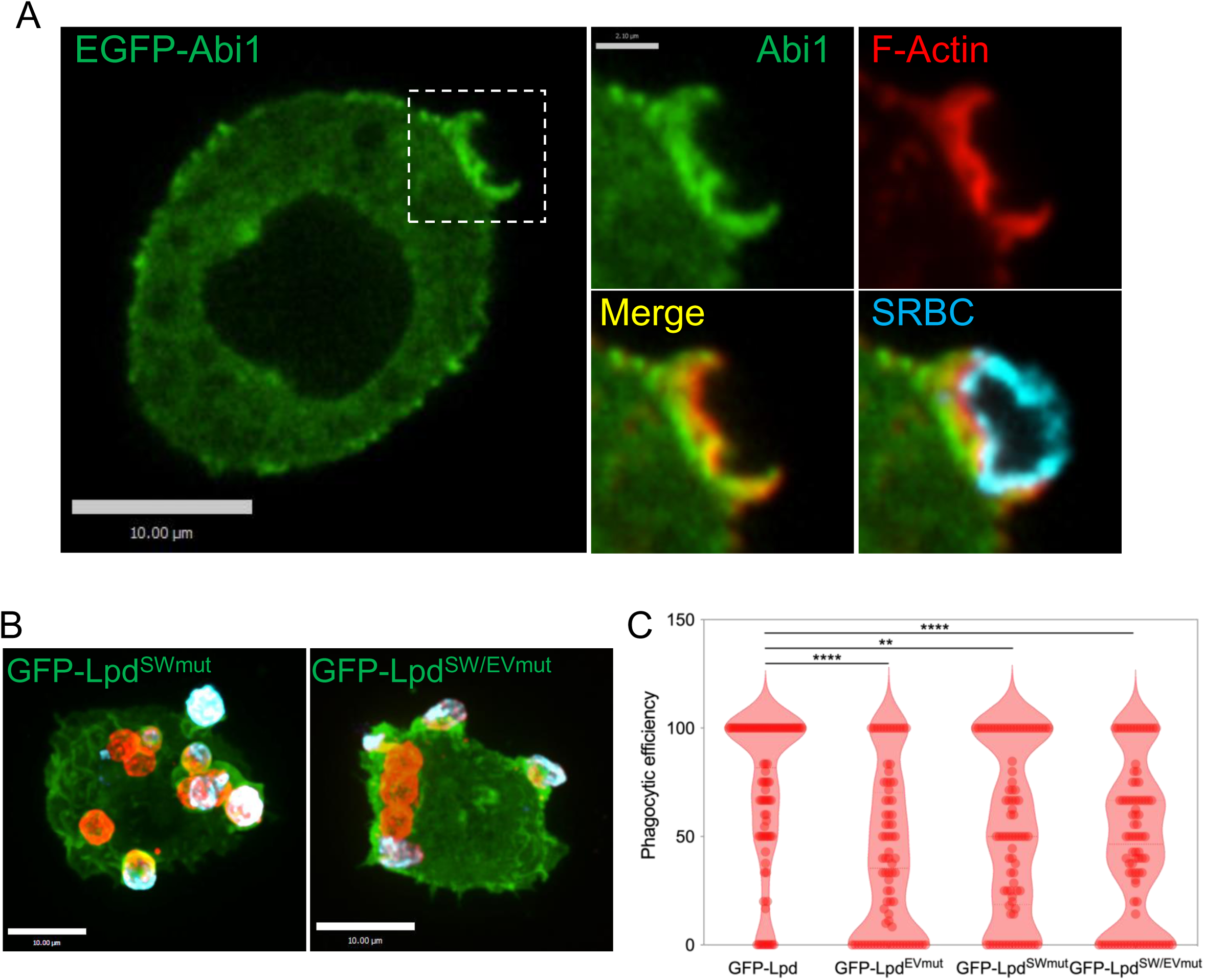
Lamellipodin-WRC interactions positively regulate phagocytosis. (A) Representative confocal slice of a RAW 264.7 macrophage expressing the WRC protein Abi1- GFP during phagocytosis and stained with phalloidin. Insets represent the area denoted by the dotted box. Similar results were obtained in 3 independent experimental repeats. (B) Representative confocal extended-focus projections of RAW 264.7 macrophages expressing Lpd^SWmut^ (left) and Lpd^SW/EVmut^ (right) after a 10-min incubation with IgG-opsonized SRBCs. Inside-outside staining was performed to differentiate between internalized/total (red) and non-internalized (blue) SRBCs. (B) Quantification of the phagocytic efficiency in RAW 264.7 macrophages overexpressing either Lpd^WT^, Lpd^EVmut^, Lpd^SWmut^, and Lpd^SW/EVmut^ . One-way ANOVA of individual phagocytic efficiency values n=86 (Lpd), n=80 (Lpd^EVmut^), n=84 (Lpd^SWmut^), n=88 (Lpd^SW/EVmut^), across N=3 independent experiments; data are means ± SEM.

**Supplemental Figure 10.**
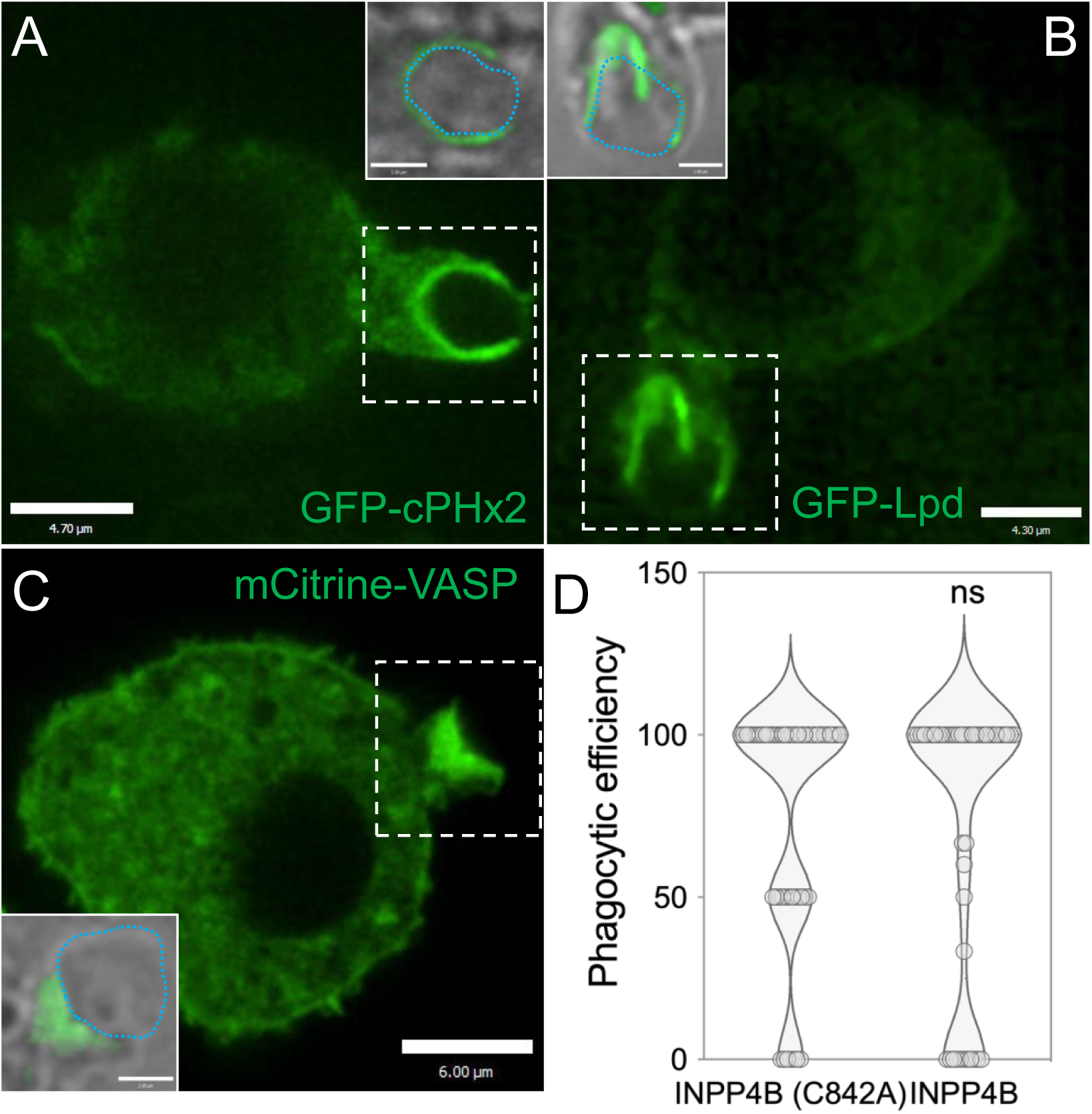
PtdIns(3,4)P_2_, Lamellipodin, and VASP Localize to the phagocytic cup during CR3-mediated phagocytosis, but PtdIns(3,4)P_2_ is dispensable for CR3-mediated phagocytosis. Representative confocal slices of a RAW 264.7 macrophage expressing GFP-cPHx3 (A), GFP-Lpd (B), and mCitrine-VASP (C) during CR3-mediated phagocytosis. Insets represent the area denoted by the dotted box showing bound SRBCs. Similar results were obtained in 3 independent experimental repeats. (D) Quantification of the phagocytic efficiency in BFP- INPP4B(C842A)-CAAX and BFP-INPP4B-CAAX-expressing RAW 264.7 macrophages. Unpaired *t*- test of individual phagocytic efficiency values across N=3 independent experiments; data are mean ± SEM.

**Supplemental Video 1.**
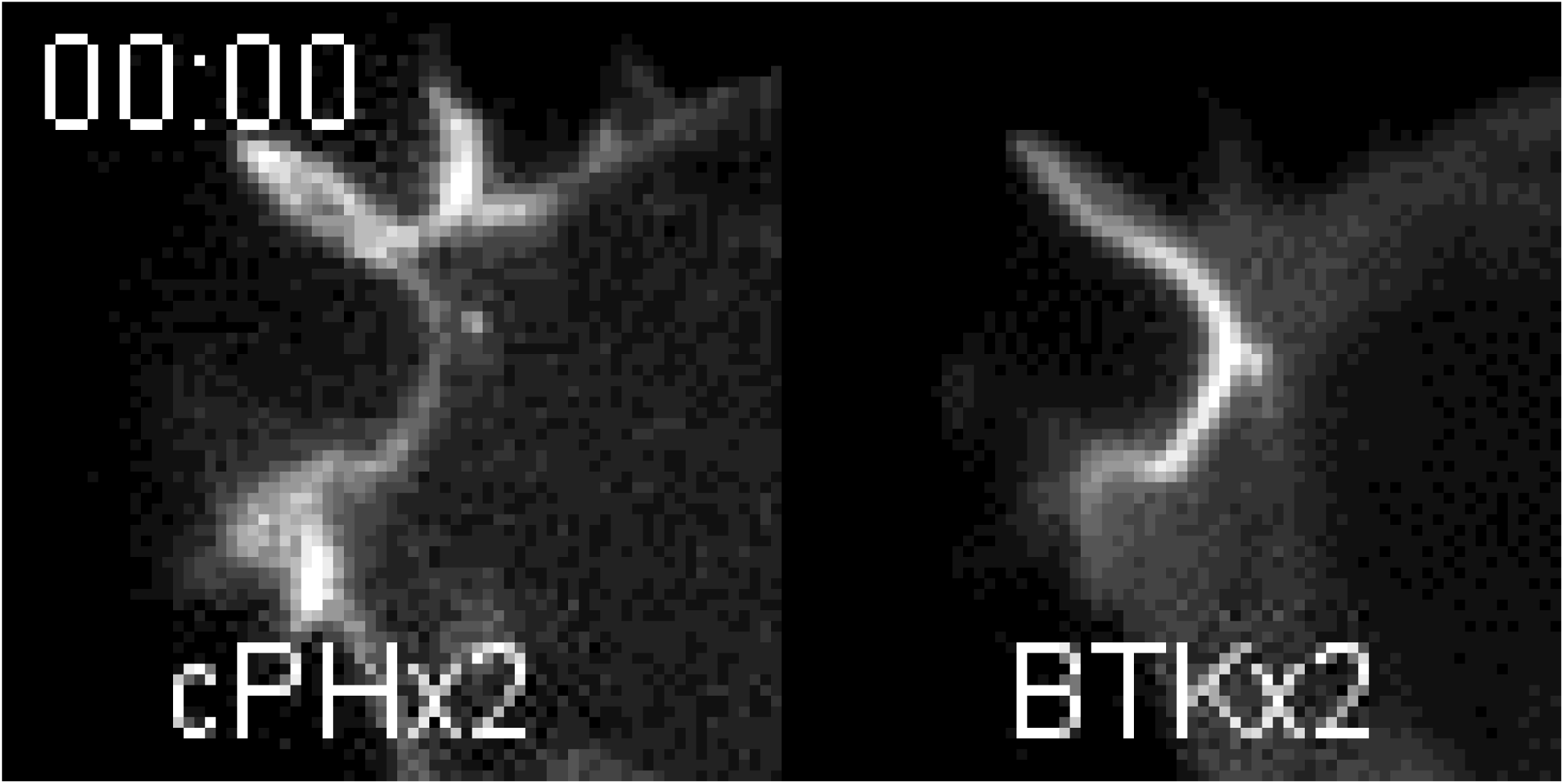
Dynamics of PtdIns(3,4)P_2_ and PtdIns(3,4,5)P_3_ during phagocytosis. RAW 264.7 macrophages co-expressing the cPHx2(left) and PH-BTKx2 (right) during phagocytosis.

**Supplemental Video 2.**
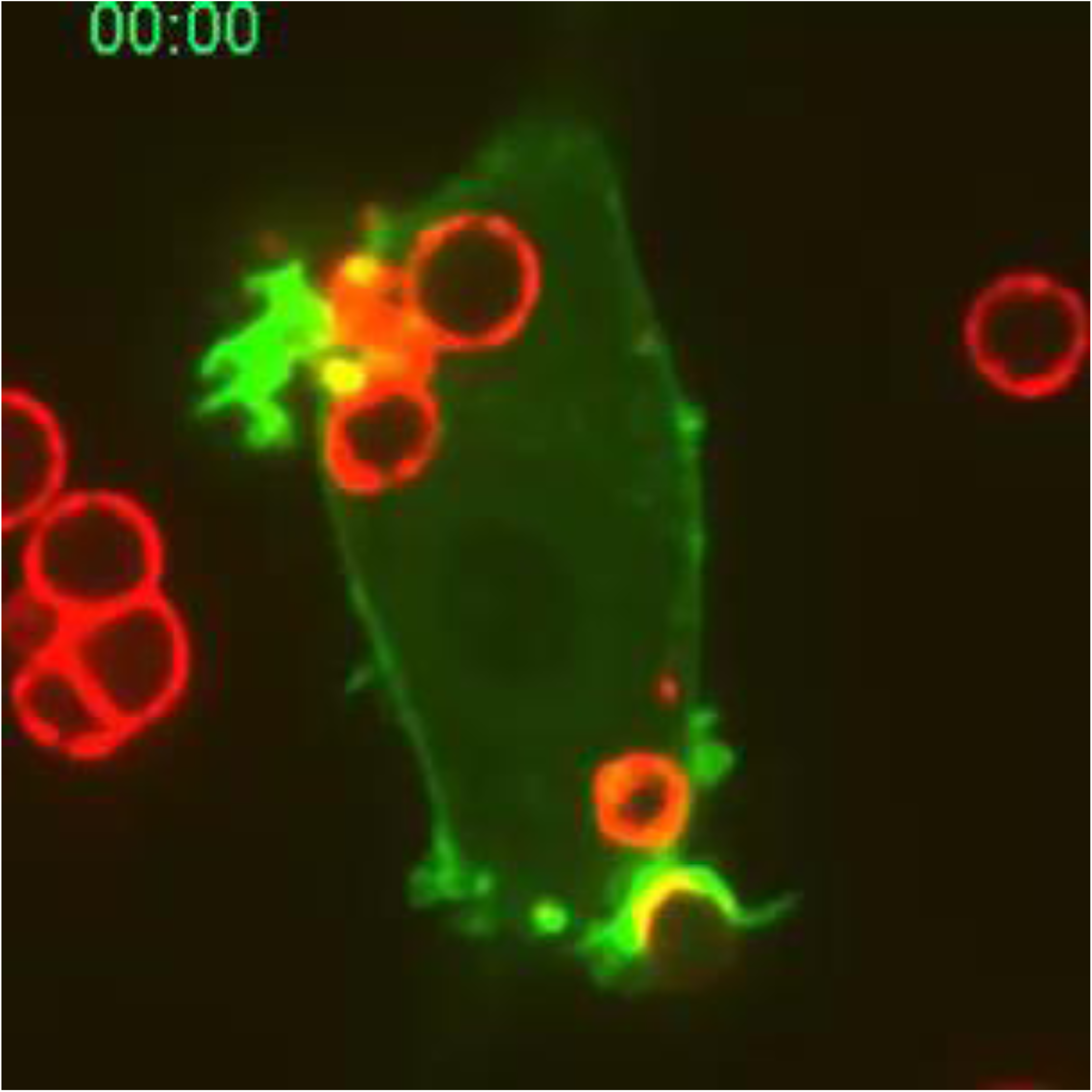
Actin dynamics during phagocytosis. Actin dynamics in RAW 264.7 macrophages expressing the INPP4B(C842A)-CAAX construct. Similar results were observed in at least 3 independent experiments.

**Supplemental Video 3.**
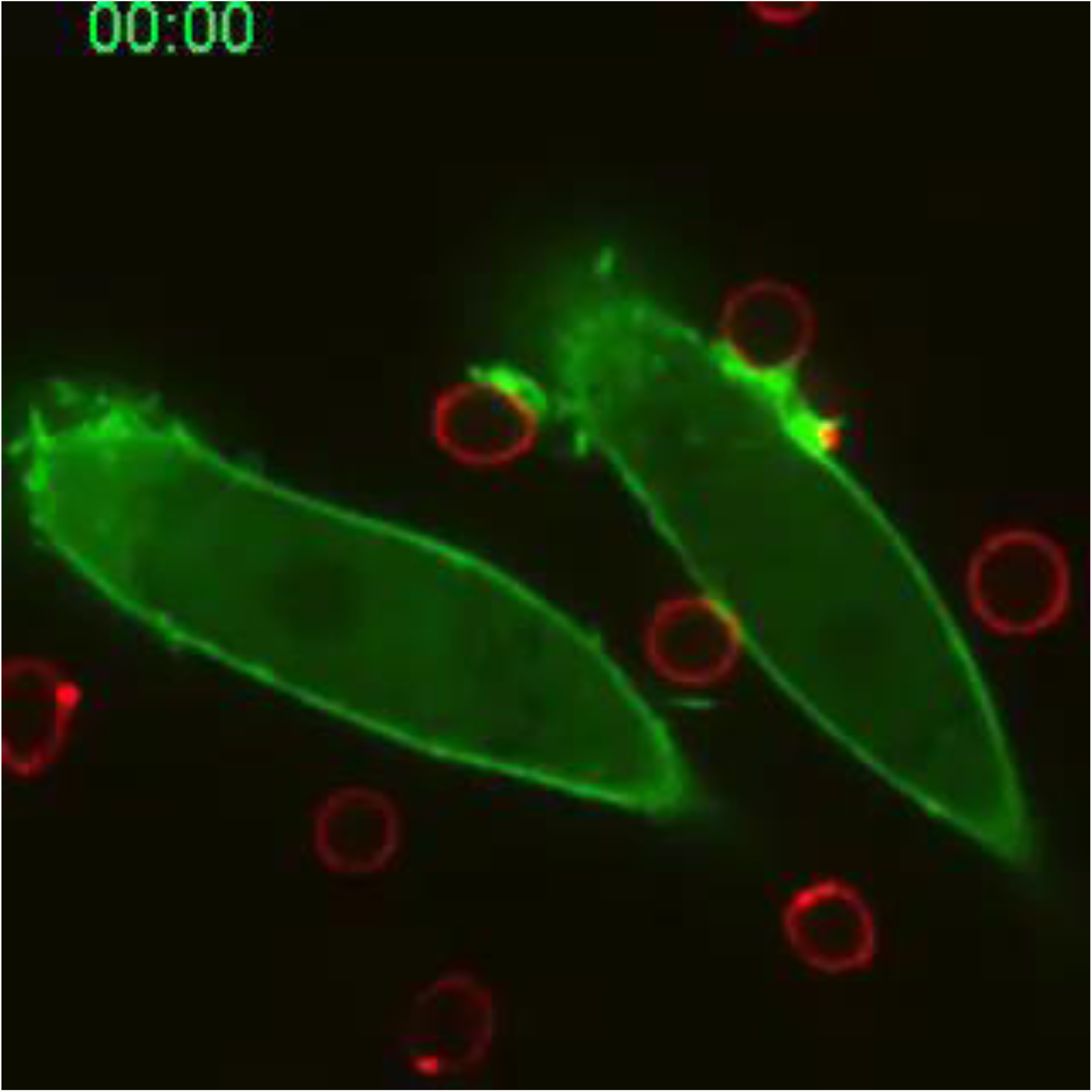
Actin dynamics in PtdIns(3,4)P_2_-depleted macrophages during phagocytosis. Actin dynamics in RAW 264.7 macrophages expressing INPP4B-CAAX construct. Similar results were observed in at least 3 separate experiments.

## Response to previous review

**Reviewer #1** (Evidence, reproducibility and clarity (Required)):

The authors use newly available probes to show that the phosphoinositide PI(3,4)P2 plays a previously undescribed role in FcgammaR-mediated phagocytosis. Using RAW macrophages, they show that PI(3,4)P2 is enriched at the plasma membrane and also at phagocytic cups internalizing IgG-opsonized sheep red blood cells. Pharmacological inhibition using wortmannin, and also expression of membrane-targeted INPP4B phosphatase showed that PI(3,4)P2 production depends on PI3 kinase activity. Further experiments using selective inhibitors showed that PI(3,4)5P2 is mainly derived by dephosphorylation of PI(3,4,5)P3, likely by multiple phosphatases such as SHIP1 or OCRL. Depletion of PI(3,4)P2 at the plasma membrane by INPP4B also resulted in strongly decreased internalization of red blood cells, although their attachment to macrophages seemed unaltered, pointing to defects in particle engulfment.

The authors then tested the potential role of lamellipodin, one of the few known PI(3,4)P2 specific effectors. Lamellipodin was found to be enriched at phagocytic cups, and this enrichment was shown to be dependent on the presence of PI(3,4)P2, by targeting of INPP4B to the plasma membrane. Macrophages depleted of lamellipodin by shRNA treatment showed reduced phagocytic efficiency and also aberrant phagocytic cup formation. As VASP is a known binding partner of lamellipodin and involved in actin polymerization, the authors next tested its potential involvement. Overexpression experiments showed that VASP colocalizes with lamellipodin at phagocytic cups. Sequestering of VASP at mitochondria through a respective construct containing the VASP binding site of ActA, together with a mitochondrial targeting sequence, showed that this also results in incompletely formed phagocytic cups and reduced phagocytic efficiency. Similar effects were observed upon expression of a lamellipodin construct with mutated binding sites for VASP.

Collectively, the authors propose that PI(3,4)P2 is localized produced at phagocytic cups through the sequential activity of PI3 kinase and PI5 phosphatase, that it recruits lamellipodin and its binding partner VASP, and that this cascade is necessary for proper phagocytic cup formation and closure and thus phagocytic capacity of cells.

This is an interesting study that uncovers a novel role for PI(3,4)P2 in phagocytic cup formation and closure. It is very well controlled, and the claims of the study are supported by the presented data. Statistical analysis is sound.

Major comments:

1) The localization of VASP at phagocytic cups is only shown by overexpression of constructs. Endogenous staining of VASP should support this finding.

We agree with the reviewer that localization of the endogenous VASP would strengthen our conclusions. We have therefore performed the suggested experiments and in the revised manuscript include a new panel (E) in the revised Figure 6 showing immunostaining of endogenous VASP during phagocytosis. The result confirms the localization of the GFP-chimeric protein.

2) It is unclear whether the roles of PI(3,4)P2, lamellipodin, and VASP are restricted to FcgammaR-mediated phagocytosis. Their potential involvement in CR3-mediated phagocytosis should be discussed or addressed in a basic set of experiments.

In the revised manuscript we have extended our original observations to analyze also CR3- mediated phagocytosis, as recommended by the reviewer. A new supplemental figure (Supplemental Figure 10) now documents that PtdIns(3,4)P_2_ is also accumulated and Lamellipodin and VASP are recruited to the phagocytic cup during CR3-mediated phagocytosis. These results imply that the role of this lipid and its effectors extend to other modes of phagocytosis. These new observations are discussed on Page 14 of the revised manuscript.

Minor comments:

1) A very recent study (Körber and Faix, EJCB, 2022) describes the role of VASP in macroendocytosis in Dictyostelium. Specifically, VASP is found to be important for proper cup closure. The results are of direct importance to the current study and should be cited accordingly.

Thank you for bringing this study to our attention. We now discuss the findings of Körber & Faix on Page 9 of the revised text.

2) direct labelling of the figures would be helpful in assessing the manuscript

To facilitate re-assessment of the paper, we have added the Figure numbers directly to the individual figures in the manuscript as suggested.

Reviewer #1 (Significance (Required)):

This study highlights the role of an underappreciated phospholipid in phagocytosis. It also describes for the first time a role for lamellipodin in formation of phagocytic cups and confirms the recent finding that also VASP is necessary for phagocytic cup closure.

The paper should be of interest to researchers working on host-pathogen interaction, regulation of the actin cytoskeleton, and also to the general cell biological community

Revieweŕs expertise:

Actin regulation
Microtubule-based transport
Adhesion, migration, invasion
Phagocytosis

We thank the reviewers for his/her comments and suggestions that have clearly improved the manuscript.

**Reviewer #2** (Evidence, reproducibility and clarity (Required)):

Review of Montano-Rendon et al: ’PI(3,4)P2, Lamellipodin and VASP coordinate cytoskeletal remodeling during phagocytic cup formation in macrophages’

The authors employ biosensors for PI(3,4)P2 in RAW 264.7 macrophages to identify localized pools of PIP2 that were sensitive to INPP4B and wortmannin (Fig1). The biosensors for PIP2 are enriched on the forming phagocytic cup (Fig2, movies) in these macrophage cells. Inhibitors for PI3K blocked the recruitment of this biosensor to the membrane.

Overall, the data are clear with the exceptions noted below. Krause et al (Dev Cell 2004) published a manuscript looking at PIP2, Lpd, and VASP in non-macrophage cells (fibroblasts, HeLa, etc…) where the influence of PI(3,4)P2 and these proteins was found to regulate actin and lamellipodial membrane extensions. This study also implicated Lpd protein coordinated actin networks in the docking of pathogens such as Vaccinia virus and EPEC bacteria. Given the additional reports of these proteins participating in dorsal ruffling (Michael et al Curr Bio 2010) and invasion (Carmona et al Oncogene 2016), it comes as no surprise that they participate in phagophore formation and phagocytosis. These studies are referenced, but having this in mind does diminish the novelty of implicating Lpd and VASP in the phagocytic process, though it seems to be the first time this machinery was directly implicated in macrophage cells.

We would like to point out that docking of viruses or dorsal ruffling are very different biological processes from phagocytosis and that the common involvement of Lamellipodin in these very disparate processes does not, in our view, detract from the novelty of our studies.

Specific Comments

Although the images and movies graphically demonstrate a PI(3,4)P_2_ enrichment on phagocytic structures, the authors could provide some additional images that include fluorescently tagged phagocytic cargo such as the erythrocytes used. The addition of a fluorescent marker or phase image would be especially beneficial in the experiments where a lack of cPHx-biosensor recruitment is seen to the docked phagocytic cargo.

We thank the reviewer for this suggestion. In the revised manuscript figures now include micrographs of the fluorescently labelled particles or phase-contrast images where appropriate.

Otherwise, readers are left with the impression that perturbations such as INPP4B compromise docking and phagocytic cup formation altogether (Fig 2C)- which is perhaps the authors point? Make this clear?

We apologize for the ambiguity of the former version of the manuscript. Initially, we noticed that particle engulfment −which is what we believe the reviewer means by “cup formation”− was the main defect in INPP4B-CaaX expressing cells. However, since the reviewer raised the possibility, we have gone back and re-analyzed the data and found that cells expressing the INPP4B-CaaX also have a small (∼35%) decrease in particle engagement/binding (Reviewer uses the term “docking”). This suggests that the plasmalemmal pool of PtdIns(3,4)P_2_ in resting cells supports the actin dynamics at the cell surface which allows the RAW cells to survey their immediate environment and thereby contact more potential prey. This new finding is included in and discussed the revised manuscript. We thank the reviewer for prompting us to consider this alternative mechanism.

There has already been an implication for PI3K in the phagocytic process, perhaps verifying that initial formation/membrane extension stages of phagocytosis are impacted by targeting the D-4 position of PIP2 would be of interest?

PtdIns(4,5)P_2_ is well known to be essential for actin polymerization and is increased transiently at the sites of phagocytosis (Botelho et al., 2000 *J Cell Biol*; Scott et al., 2005 *J Cell Biol;* Fairn et al., 2009 *J Cell Biol*.). It is not clear whether the reviewer is curious about the possible consequences of converting PtdIns(4,5)P_2_ to PtdIns5P prior to activation of PI3K. Whether PtdIns5P itself has biological activity is a subject of debate and, to our knowledge, its existence has not been documented at sites of phagocytosis. It is also unclear whether PtdIns5P would serve as an effective substrate for PI3K and, if so whether the putative product, PtdIns(3,5)P_2_ that is normally found in endomembranes, would be functionally relevant.

Depletion of PI(3,4)P2 through the expression of the INPP4B phosphatase demonstrated a reduction in phagocytic uptake of red blood cells (Fig4). The readout for this assay relied upon what appears to be differential labeling of phagocytosed red blood cells, though there are examples of cargo that is supposedly inside the macrophages labeled in green? Perhaps the authors can reconcile this and make the methods more clear for this approach?

Thank you for this comment; we apologize if the original text was unclear in this regard. In the revised manuscript a detailed description of the staining protocol we used to distinguish inside from outside particles is now included in the Methods section. It is also worth pointing out that the green-only SRBCs in the INPP4B-CaaX panel in Figure 3 indicate targets that were fully internalized by those RAW macrophages <u>not</u> expressing BFP-INPP4B-CAAX (see image below)

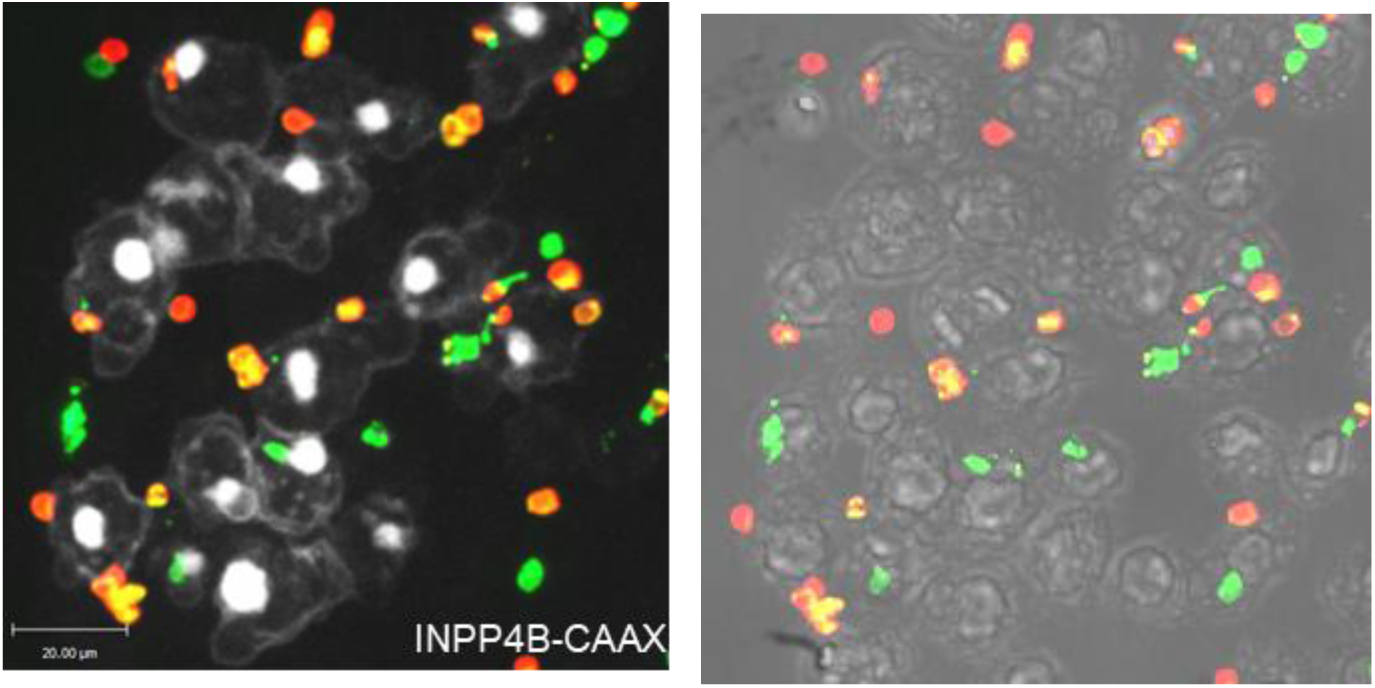

Fig4 demonstrates the PI(3,4)P2 dependent recruitment of Lamellipodin (Lpd) to the phagocytic cup, which is clear. Lpd is found to be necessary for effective phagocytic uptake in Fig 5. There is no blotting/qPCR data for the verification of Lpd knockdown shown?

RAW macrophages and other macrophage cell lines are rather refractory to transfection, resulting in only a minor (10-20%) fraction of the cells expressing transfected constructs. For this reason, immunoblotting or qPCR analyses of the entire population yield misleading results, not reflective of the comparatively small transfected sub-population of cells. To overcome this limitation, we co-transfected the shRNA-containing plasmid with a smaller amount of a plasmid containing a fluorescent protein used to identify transfectants visually (a 5:1 ratio of shRNA:EGFP). By using a 5:1 ratio of the plasmids we ensured that cells expressing the fluorescent protein had a high likelihood of also expressing the shRNA. In this manner, the Lpd- depleted cells could be scored separately from the untransfected, wild-type cells following immunostaining (Supplemental Figure 5). Note that some immunostaining persisted in the Lpd- silenced cells, in all likelihood because some of the antibody binding is nonspecific, as is commonly seen in immunostaining. Nevertheless, the data indicate that substantial silencing of Lpd is achieved when transfecting the shRNA.

The authors demonstrate a co-localization of Lpd/VASP proteins at the phagocytic cup of these macrophages in Fig 6 and sequester VASP protein to the mitochondria with some ActA derived fusion proteins to functionally block phagocytosis. The functional interaction of Lpd/VASP is further explored with experiments utilizing Ena/VASP mutants in Fig7, demonstrating a dependence on this interaction to promote phagocytic uptake.

Reviewer #2 (Significance (Required)):

see above

**Reviewer #3** (Evidence, reproducibility and clarity (Required)):

In this study, Montano-Rendon and colleagues address the role of phosphatidylinositol (3,4)- bisphosphate in phagocytosis by RAW macrophages. Using small molecule inhibitors, they show that dephosphorylation of PI(3,4,5)P3 is the main source of PI(3,4)P2 in phagocytosis. Using an elegant approach based on overexpression of a PI(3,4)P2-specific phosphatase, they show that the selective depletion of PI(3,4)P2 impairs phagosome formation. Moreover, they identify two PI(3,4)P2 interacting proteins involved in phagocytosis: lamellipodin and VASP. They show that shRNA silencing of lamellipodin arrests phagocytosis, as well as mistargeting of VASP to mitochondria by a fusion protein. Overall, this is a high-quality study, well designed and written. I hence support publication, and only have a few relatively minor comments that the authors should consider as I believe it would improve the quality of the manuscript.

The role of PI(3,4)P_2_ in the actin organisation in phagocytosis has been shown previously in various studies, see for example PMID: 16418223, 27806292 and review 32296634. In these studies, different mechanisms have been proposed of how PI(3,4)P2 affects the cytoskeleton and phagocytic process. It would be good to discuss how the findings with lamellipodin and VASP relate to these previously described mechanisms.

We now include and discuss the references recommended by the reviewer to highlight that the importance of PtdIns(3,4)P_2_ extends to dendritic cells and HL60 neutrophils.

In figure 6, a role for VASP in phagocytosis is shown by mistargeting it to mitochondria using a fusion protein consisting of a VASP binding region and a mitochondrial targeting motif. While this is an elegant approach, I wonder why not simply shRNA is used, similar to lamellipodin?

We decided to use this approach because macrophages (including RAW cells) express other members of the Ena/VASP family of proteins such as EVL (Coppolino et al., 2001 *J Cell Sci)* that could potentially substitute for VASP; simultaneously silencing multiple, distinct members of the Ena/VASP family poses an experimental challenge. Moreover, in our experience introducing siRNA into RAW cells, even when using electroporation, is often insufficient to generate robust silencing of certain genes (e.g. Levin-Konigsberg, et al., 2019 *Nature Cell Biology*). Thus, we took advantage of the robust, more globally effective ActA-based molecular tool. To demonstrate its effectiveness, we now include a new Supplemental figure (Supplemental Figure 6, reproduced below) using immunostaining that shows how virtually all of the endogenous VASP is sequestered to the surface of mitochondria when the MITO-FP4 is expressed.

**Supplemental Figure 6.**
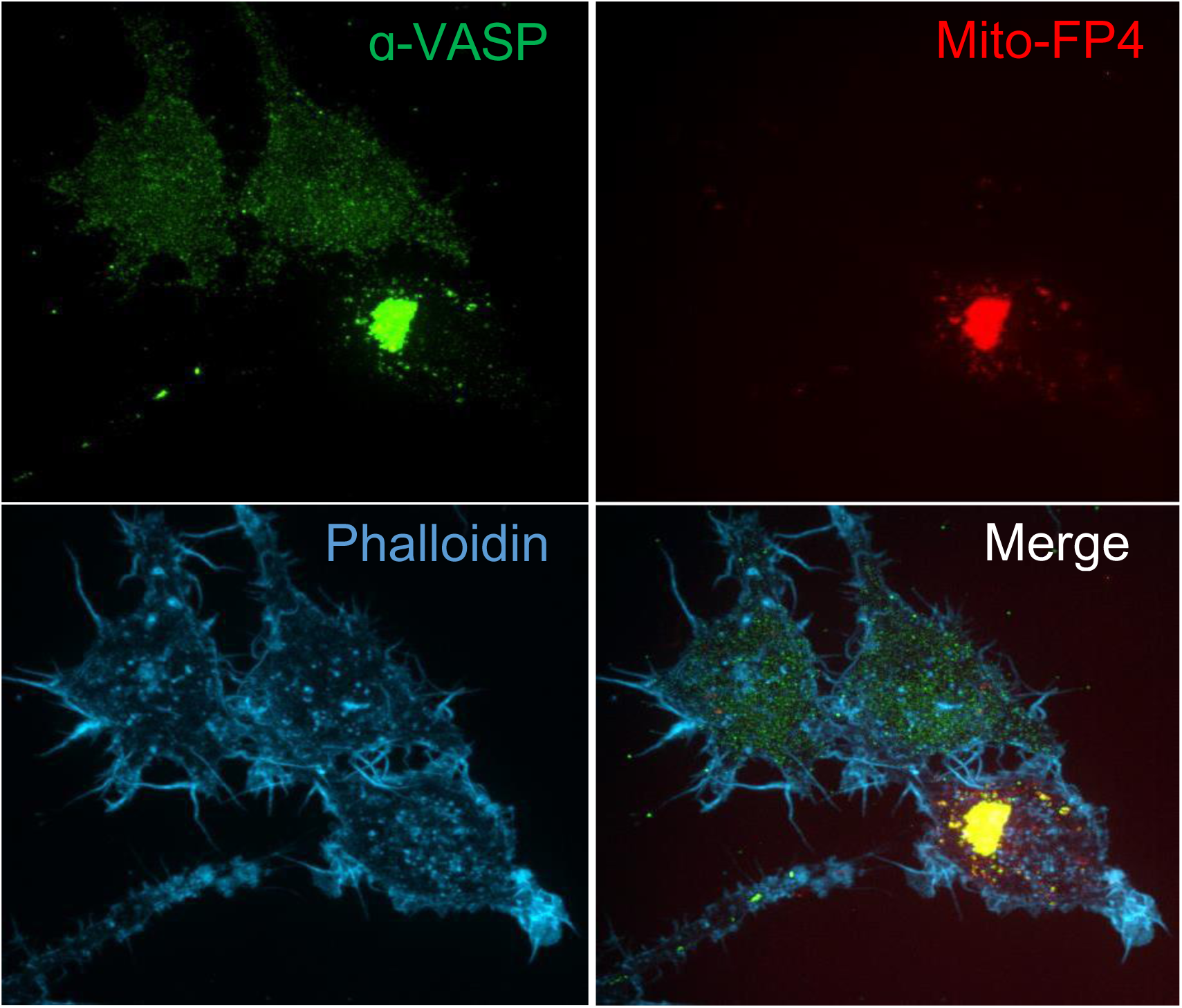
MITO-FP4 targets endogenous VASP to the Mitochondria.

In figure 3A: How was the inside-outside staining performed? I cannot find this information in the Methods.

We apologize for the omission. The inside/outside staining protocol is now detailed in the Methods section of the manuscript.

Figures are overall good quality. However, in figure 1,2, and 4 individual cells are shown in the graphs, whereas figures 3, 5, 6 and 7, and the supplementary figures only show averages with bar graphs. Please change these graphs to all show individual cells, as this will allow to see the variation among cells.

Thank you for the suggestion. The graphs have been modified to violin plots to show the variation and distribution of results amongst the individual cells and experiments.

Reviewer #3 (Significance (Required)):

see above

